# Systemic circulating microRNA landscape in Lynch syndrome

**DOI:** 10.1101/2022.03.10.483718

**Authors:** Tero Sievänen, Tia-Marje Korhonen, Tiina Jokela, Maarit Ahtiainen, Laura Lahtinen, Teijo Kuopio, Anna Lepistö, Elina Sillanpää, Jukka-Pekka Mecklin, Toni T. Seppälä, Eija K. Laakkonen

**Affiliations:** Gerontology Research Center and Faculty of Sport and Health Sciences, University of Jyväskylä, Jyväskylä, Finland; Department of Education and Research, Central Finland Health Care District, Jyväskylä, Finland; Department of Pathology, Central Finland Health Care District, Jyväskylä, Finland; Department of Biological and Environmental Science, University of Jyväskylä, Jyväskylä, Finland; Department of Surgery, Abdominal Center, Helsinki University Hospital, Helsinki, Finland; Institute for Molecular Medicine Finland, University of Helsinki, Helsinki, Finland; Department of Surgery, Central Finland Health Care District, Jyväskylä, Finland; Faculty of Sport and Health Sciences, University of Jyväskylä, Jyväskylä, Finland; Applied Tumor Genomics Research Program, University of Helsinki, Helsinki, Finland

**Author notes:** **Corresponding author:** Eija K. Laakkonen, University of Jyväskylä, P.O. Box 35 (VIV), 40014, Jyväskylä, Finland. Equal contribution (Toni T. Seppälä, Eija K. Laakkonen).

**Keywords:** Hereditary cancer, Lynch syndrome, microRNA, mismatch-repair, next generation sequencing

## Abstract

MicroRNAs (miRs) are non-coding RNA-molecules that regulate gene expression. Global circulating miR (c-miR) expression patterns (c-miRnome) change with carcinogenesis in various sporadic cancers. Therefore, aberrantly expressed c-miRs could have diagnostic, predictive and prognostic potential in molecular profiling of cancers. c-miR functions in carriers of inherited pathogenic mismatch-repair gene variants (*path_MMR*), also known as Lynch syndrome (LS), have remained understudied. LS cohort provides an ideal population for biomarker mining due to increased lifelong cancer risk and excessive cancer occurrence. Using high-throughput sequencing and bioinformatic approaches, we conducted an exploratory analysis to characterize systemic c-miRnomes of *path_MMR* carriers. Our discovery cohort included 81 healthy *path_MMR* carriers and 37 non-LS controls. Our analysis also included cancer cohort comprised of 13 *path_MMR* carriers with varying cancers and 24 sporadic rectal cancer patients. We showed for the first time that c-miRnome can discern healthy *path_MMR* carriers from non-LS controls but does not distinguish healthy *path_MMR* carriers from cancer patients with or without *path_MMR*. Our c-miR expression analysis combined with *in silico* tools suggest ongoing alterations of biological pathways shared in LS and sporadic carcinogenesis. We observed that these alterations can produce a c-miR signature which can be used to track oncogenic stress in cancer-free *path_MMR* carriers. Thus, c-miRs hold potential in monitoring which cancer patients would require more intensive surveillance or clinical management.

**Significance:** C-miRnome can discern between healthy persons with or without *path_MMR* but does not distinguish healthy *path_MMR* carriers from cancer patients with or without *path_MMR*, indicating an ongoing alteration of biological pathways that can be used to track oncogenic stress at cancer-free state.

## Introduction

Lynch syndrome (LS) is an inherited cancer predisposition syndrome caused by pathogenic gene variants in DNA-mismatch repair (*path_MMR*) genes *MLH1, MSH2, MSH6* or *PMS2* (1). By genetic or epigenetic silencing, deficient MMR (dMMR) significantly increases cellular mutation rates thus predisposing *path_MMR* carriers to increased cancer risk and excessive cancer occurrence (1,2). Colorectal cancer is a traditional hallmark cancer of LS that is commonly cured by surveillance, followed by modern surgical and oncological management, with over 90% 10-year overall survival (2,3). Despite the good recovery rate in first cancers, the persons at risk will develop frequently more lethal cancers still at relatively young age (4). This highlights the need for an improved molecular assessment and identification of which patients would require more intensive surveillance or clinical management.

MicroRNAs (miRs) are small (18-25 nucleotides) non-coding RNA-molecules that regulate gene expression by translational repression (5). MiRs play a role in regulation of >30% of the human genes controlling critical biological processes such as cell proliferation, cell differentiation, and apoptosis (5–7). In cancers, miRs can be regarded as tumor suppressive or oncogenic, thus resulting in downregulation or upregulation of the affected target genes, respectively (7). Compared to tissue-based miRs, circulating-miRs (c-miRs) migrate throughout the body within various body fluids and are part of active inter-tissue crosstalk (8,9). Nowadays, profiling of the global c-miR expression levels (c-miRnome) has become prevalent and miR expression can be correlated with cancer type, stage, and other clinical variables (10–13). Therefore, aberrantly expressed miRs could have diagnostic, predictive, and prognostic potential in molecular profiling and early detection of cancers.

LS cohort provides an ideal population for biomarker mining due to well-predicted cancer risk of persons under frequent surveillance. The role of miRs in LS have remained understudied even if various studies have shown that c-miR expression patterns change with carcinogenesis in various sporadic cancers. Balaguer et al. have shown that miRs can be used in tumor classification and discrimination of sporadic and hereditary tumors with microsatellite instability (14), thus highlighting the potential role of miRs as LS biomarkers. In support, Valeri et al., Liccardo et al. and Zhou et al. postulated that miRs could have functional roles in LS carcinogenesis, e.g., by targeting MMR-proteins (15,16) and various tumor-suppressor genes (17). However, these studies along with other reports have assessed miR functions in the colorectum and colorectal cancer tissues and cells as well as with microarray data *in silico* (14–19) but not in circulation.

Instead of using a targeted panel of *a priori* chosen c-miRs, it is beneficial to characterize the systemic c-miRnome of *path_MMR* carriers. This “omics-approach” provides a more comprehensive view of how c-miRs could contribute to LS pathogenesis, and plausibly pave way for future use of c-miRs in risk stratification and early detection of LS cancers. Our exploratory study compared the systemic c-miRnome of cancer-free *path_MMR* carriers with c-miRnomes of non-LS controls (discovery cohort), sporadic rectal cancer patients and *path_MMR* carriers with cancer (cancer cohort) using high-throughput sequencing and bioinformatic approaches.

## Materials and methods

### Study subjects

This study consisted of independent discovery and cancer cohorts. The discovery cohort (n=118) was composed of 81 currently cancer-free (healthy) Finnish *path_MMR* carriers and 37 non-LS controls whose c-miRnomes were sequenced. The cancer cohort (n=37) was composed of 13 *path_MMR* carriers who currently had cancer and 24 sporadic rectal cancer patients whose c-miRnomes were sequenced.

All *path_MMR* carriers were enrolled in the study at their regular colonoscopy surveillance appointments. They were also registered participants in the nationwide Finnish Lynch Syndrome Research Registry (LSRFi, www.lynchsyndrooma.fi, accessed 05/2021). The families and individuals were identified in the registry based on clinical criteria (Amsterdam and Bethesda criteria) (20,21) and subsequently through cascade testing of the families and universal testing of tumors. Adult members of LSRFi with confirmed *path_MMR* variants (classes 4 and 5 by InSiGHT criteria) (22) were eligible for the study.

Sporadic rectal cancer patients were enrolled at the time of their initial appointment for surgery at surgical clinic at the local tertiary center responsible for management of rectal cancer in the Southern Finland area (Helsinki University Central Hospital, unit of rectal surgery).

Non-LS control samples were acquired from Biobank of Eastern Finland (n=27) in 2020 or were part of the Estrogenic Regulation of Muscle Apoptosis (ERMA) cohort (n=10) consisting of healthy 47-55-years old women (23). Persons with no cancers, blood disorders, acute or chronic infectious diseases, rheumatoid arthritis and known *BRCA* or MMR-gene germline mutations were eligible for the non-LS control group. Ethnicity throughout the study population was widely white Caucasian.

### Ethics statement

Informed consent was obtained from all participants, and the Helsinki and Uusimaa Health Care District (HUS/155/2021) and Central Finland Health Care District Ethics Committee (KSSHP D# 1U/2018 and 1/2019 and KSSHP 3/2016) approved the study protocol. The study was conducted according to the guidelines of the Declaration of Helsinki.

### Sample collection

*Path_MMR* carriers’ and sporadic rectal cancer patients’ venous blood samples were drawn after surveillance colonoscopy visits at fasted state. All ERMA participants fasted overnight before blood sampling. The duration of fasting is not reported for the samples obtained through biobank (n=27). Samples were taken from antecubital vein to standard serum tubes (455092, Greiner). To separate serum, the whole blood samples were allowed to clot for 30 minutes at room temperature, centrifuged at 1,800 *g* for 10 min and aliquoted.

### Small-RNA isolation and quality evaluation

c-miR isolations from blood serum were carried out using affinity column-based miRNeasy Serum/Plasma Advanced Kit (217204, Qiagen) according to the manufacturer’s instructions. Briefly, 0.5 ml of thawed serum was used to isolate miRs. All the required solutions were added in amounts recommended by the manufacturer. Cel-miR-39 miR mimic (MS00019789, Qiagen) was added to each sample to serve as a spike-in control for monitoring the miR purification and amplification. Phase separation centrifugation was executed in 12,000 *g* for 3 min at room temperature (Heraeus, Biofuge Pico and Fresco 17, ThermoFisher) and rest of the centrifugations were performed at 16,000 *g* whenever a range of 8,000–20,000 *g* was recommended. C-miRs were eluted to nuclease-free water. Prior to the library preparation, RNA quality and recovery were checked by RT-qPCR (CFX96-RT-qPCR, Bio-Rad) according to manufacturer’s protocol (MiScript Primer assays and II RT kit for cDNA synthesis and MiScript SYBR Green PCR Kit for RT-qPCR, 218161, Qiagen) from which the recovery of cel-miR-39 spike-in control was confirmed.

### Small-RNA library preparation and sequencing

Small-RNA Library preparations were executed with QIAseq miRNA Library Preparation Kit (1103679, Qiagen) according to the manufacturer’s instructions using multiplexing adapters. Briefly, the small RNA fractions were first ligated to sequencing adapters from both 5’ and 3’ ends, reverse transcribed into cDNA using UMI-assigning primers and purified using magnetic beads. A universal indexing sequence was also added in the reverse transcription step, thus allowing samples to be distinguished from each other. The samples were then amplified with standard thermocycler (Eppendorf), purified and eluted into nuclease-free water. Quality assessment of the libraries was completed with TapeStation 4200 (Agilent). The library sample concentrations were measured with Qubit fluorometer (Invitrogen), quantified, diluted and pooled into a single mixture in equal amounts (1.8 pM per sample) prior to sequencing. Sequencing of the small-RNA libraries were done with NextSeq 500 (Illumina) using NextSeq 500/550 High Output Kit v. 2.5 with 75 cycles (15057934, Illumina) to produce 75-base pair single-end reads with aimed mean sequencing depth of >5 M reads per sample as recommended by the manufacturer (Qiagen).

### Raw data processing and alignment

Sequencing output data was converted to FASTQ-format using bcl2fastq software (v.2.20, Illumina, USA). FastQC was used for quality controls (24). The QIAseq sequencing adapters were trimmed from the 3’ end of the reads with FastX-toolkit (25) using default parameters with minimum alignment length -M 19. Only clipped reads >20 bp in length were selected for downstream analysis. After adapter clipping, the reads were trimmed to 22 bp to enrich miR-sequences and then quality filtered with FastX-toolkit. Only high-quality reads (Phred score >25) were selected for alignment to reference genome. Before alignment, all the four sample lanes were merged to obtain the overall sample read count and to ensure better mapping quality. Samples that had <1 M reads were excluded from the analyses. Subsequently, the pre-processed reads were mapped to human mature miR-genome (miRbase v.22) (26) with Bowtie alignment tool for single end data with v-mode and best strata parameters (27). Only uniquely mapped miR-reads were selected for differential expression (DE) analysis.

### Differential expression analysis

DE analyses from raw c-miR counts were based on statistical procedures of EdgeR (28) and DESeq2 (29) packages and conducted in R-studio (v. 3.6.3) (30) (Supplementary file 3). Briefly, DE analyses were performed on c-miR raw read count matrices after the low expressed genes were filtered out and normalized with the median of ratios method in DESeq2. C-miRs that had more than 1 count per million in 70% of the samples in a group were selected for DE analyses. Filtered and normalized c-miR counts were used to set up a design matrix in DESeq2 that adjusted for sex and potential batch effect. Benjamini-Hochberg procedure in DESeq2 was used to correct for multiple testing. C-miRs that had a false discovery rate (FDR) <0.05 were considered DE.

### Dimension reduction analysis

Dimension reduction of the DESeq2-normalized data was conducted using the t-distributed stochastic neighbor embedding (t-SNE) method, which is a non-linear and unsupervised technique to simplify high dimensional data for visualization in low-dimensional space (31). t-SNE analysis was performed to identify and visualize possible clustering of subpopulations within the dataset. Rtsne package in R-studio was used with output dimensionality set to 2, perplexity set to 35 and theta set to 0.5.

### Target gene prediction and pathway analysis

Putative miR-target gene prediction was performed using mirWalk tool that utilizes a random-forest-based approach, an ensemble learning method based on multiple decision trees, to predict target genes (32,33). Only the predicted miR-target genes targeting 3’ untranslated region with experimental validation from miRTarBase (34) and which were included and verified in mirDB (35) and TargetScan (36) databases were selected for downstream gene set enrichment analysis (GSEA) (37). GSEA of gene ontology biological processes (GO: BP) and Kyoto Encyclopedia of Genes and Genomes (KEGG) (38) pathways were also conducted with mirWalk. MirWalk provides a standard enrichment analysis based on hypergeometric tests. GO and KEGG terms with FDR-corrected p-values of <0.05 were considered enriched. Cancer Gene Census of the Catalogue of Somatic Mutations in Cancer (COSMIC-CGC) (39) project database were used for target gene investigation.

### Statistical analysis

Data regarding study subjects are presented using means and standard deviations. DE-analyses were based on statistical procedures of DESeq2 package accounting for normalization and exclusion of outliers. Mann-Whitney U-test was used in the validation analysis (Supplementary file 1). Pearson correlation was used to compare gene fold correlation between the discovery and validation cohorts (Supplementary file 1). In all analyses, p-values or FDR <0.05 were considered to indicate statistical significance.

### Data availability

The data generated in this study can be accessed by contacting the corresponding author.

## Results

### A pool of 228 c-miRs is shared between the discovery and cancer cohorts

Descriptive characteristics of study subjects in the discovery cohort and cancer cohort are presented in Table 1.

**Table 1.** Descriptive characteristics of study subjects in the discovery cohort and cancer cohort.

| Variable | Discovery cohort |  | Cancer cohort |  |
| --- | --- | --- | --- | --- |
|  | *Path_MMR*,<br>healthy | non-LS,<br>healthy | *Path_MMR*,<br>cancer | Sporadic rectal cancer<br>patients |
| **N** | 81 | 37 | 13 | 24 |
| **Sex (N (%))** |  |  |  |  |
| Male | 40 (49.4) | 18 (48.6) | 10 (76.9) | 10 (41.6) |
| Female | 41 (50.6) | 19 (51.4) | 3 (23.1) | 14 (58.4) |
| **Age, years (mean± SD)** | 59.5 (10.7) | 54.9 (10.7) | 60.7 (15.3) | 69.8 (9.9) |
| **Body mass index, kg/m<sup>2</sup> (mean± SD)<sup>#</sup>** | 27.3 (5.7) | 28.0 (6.2) | 28.2 (3.4) | 27.6 (6.3) |
| ***Path_MMR* (N (%))** |  |  |  |  |
| *MLH1* | 50 (61.7) | - | 8 (61.5) | - |
| *MSH2* | 17 (21.0) | - | 2 (15.4) | - |
| *MSH6* | 12 (14.8) | - | 3 (23.1) | - |
| *PMS2* | 2 (2.5) | - | 0 (0.0) | - |
| **Previous cancers (N (%))** |  |  |  |  |
| Yes | 42 (51.9) | - | 10 (76.9) | - |
| No | 39 (48.1) | - | 3 (23.1) | - |
| **Cancer type (N (%))** |  |  |  |  |
| Colorectal cancer | - | - | 5 (38.5) | - |
| Prostate cancer | - | - | 3 (23.0) | - |
| Other cancer | - | - | 5 (38.5) | - |
| Rectal cancer | - | - | - | 24 (100.0) |

<sup>#</sup> Missing data: Discovery cohort, n=12 in *path\_MMR* carriers; Cancer cohort, n=3 in *path\_MMR* carriers. <sup>a</sup>Other cancer include esophageal cancer, n=1; spinocellular cancer, n=1; glioblastoma, n=1; gastric cancer, n=1 and thymic cancer, n=1.

Human genome encodes approximately 2,600 mature miRs (miRbase, v.22) (26). To inspect the systemic c-miR content in the discovery and cancer cohorts, we performed small-RNA sequencing experiment to characterize the serum c-miRnomes. We identified a total of 1349 distinct c-miRs in three separate sequencing runs with an average sequencing depth of 3.2M reads per sample (Supplementary file 1, Table S1 & Supplementary file 2). After processing of raw data and filtering of low expressed c-miRs, 228 c-miRs common to both cohorts were identified (Supplementary file 1, Fig. S1).

The most highly expressed c-miRs among *path_MMR* carriers with or without cancer were hsa-let-7a-5p, hsa-let-7b-5p, hsa-miR-122-5p, hsa-miR-16-5p and hsa-mir-223-3p (Supplementary file 1, Fig. S2). The most abundant c-miRs in non-LS control group were the same as in *path_MMR* carriers with or without cancer (Supplementary file 1, Fig. S3). Among sporadic rectal cancer group, the top c-miRs were otherwise the same except hsa-miR-451a replaced hsa-miR-122-5p (Supplementary file 1, Fig. S4). All these top c-miRs in total accounted for approximately 50% of all c-miR counts in all cohorts, thus displaying major overrepresentation that could have possibly affected the c-miR pool size. In summary, our sequencing analysis provided moderate coverage of c-miRnomes in LS.

### Healthy *path_MMR* carriers have a c-miRnome that differs from non-LS controls but resembles the c-miRnomes of patients with sporadic or hereditary cancer

The phenotype and cancer risk spectrum vary within LS cohort, e.g., due to *path_MMR* variant and sex (1). As our discovery cohort consisted of males and females with all *path_MMR* variants included, we first explored whether these traits influenced c-miR expression in healthy *path_MMR* carriers. We used the pool of identified 228 c-miRs to form the count matrix for all DE-analyses (Supplementary file 3). Hsa-miR-206 and hsa-miR-223-5p were observed downregulated in males compared to females and thus sex was added as a covariate to further analyses (Supplementary file 3). We did not find DE c-miRs when *path_MMR* variants were compared to each other or when *path_MLH1* carriers were compared to all other *path_MMR* variants combined (Supplementary file 3). These results show that different *path_MMR* variants do not cause heterogeneity that would generate a recognizable c-miR profile, thus suggesting a shared systemic response common to all *path_MMR* variants. Furthermore, we also tested if the c-miR expression profile is altered in persons who had had cancer or multiple cancers previously, but we did not find significant differences (Supplementary file 3).

Alterations in the immune cell abundance of normal colorectal mucosa in cancer-free *path_MMR* carriers separate them from those with cancer (40). To see whether we can identify a LS-specific c-miR signature, our primary objective was to characterize systemic c-miRnome of healthy *path_MMR* carriers, which has not been done previously. We thus performed DE-analysis within the discovery cohort and RT-qPCR validation analysis within similar but independent validation cohort (results in Supplementary file 1) to compare healthy *path_MMR* carriers to healthy non-LS controls. In DE-analysis, we found 40 out of 228 c-miRs to display aberrant expression in healthy *path_MMR* carriers (Table 2). Of them, 15 were upregulated and 25 downregulated in *path_MMR* carriers compared to non-LS controls, but the fold changes remained low varying from minimum of -0.88 to maximum of 1.25 (Fig. 1A). Hsa-miR-155-5p, hsa-let-7c-5p and -let-7e-5p and -122b-3p had the most significant upregulation within healthy *path_MMR* carriers (Table 2). Of the downregulated c-miRs, hsa-miR-15a-5p was the most significantly downregulated followed by hsa-miR-185-5p, -320a-3p and -186-5p, respectively (Table 2). Overall, aberrant expression of multiple c-miRs in healthy *path_MMR* carriers might indicate that some systemic alterations in c-miR-mediated regulation of biological pathways associated with dMMR may be ongoing even at cancer-free state in *path_MMR* carriers.

**Table 2.**
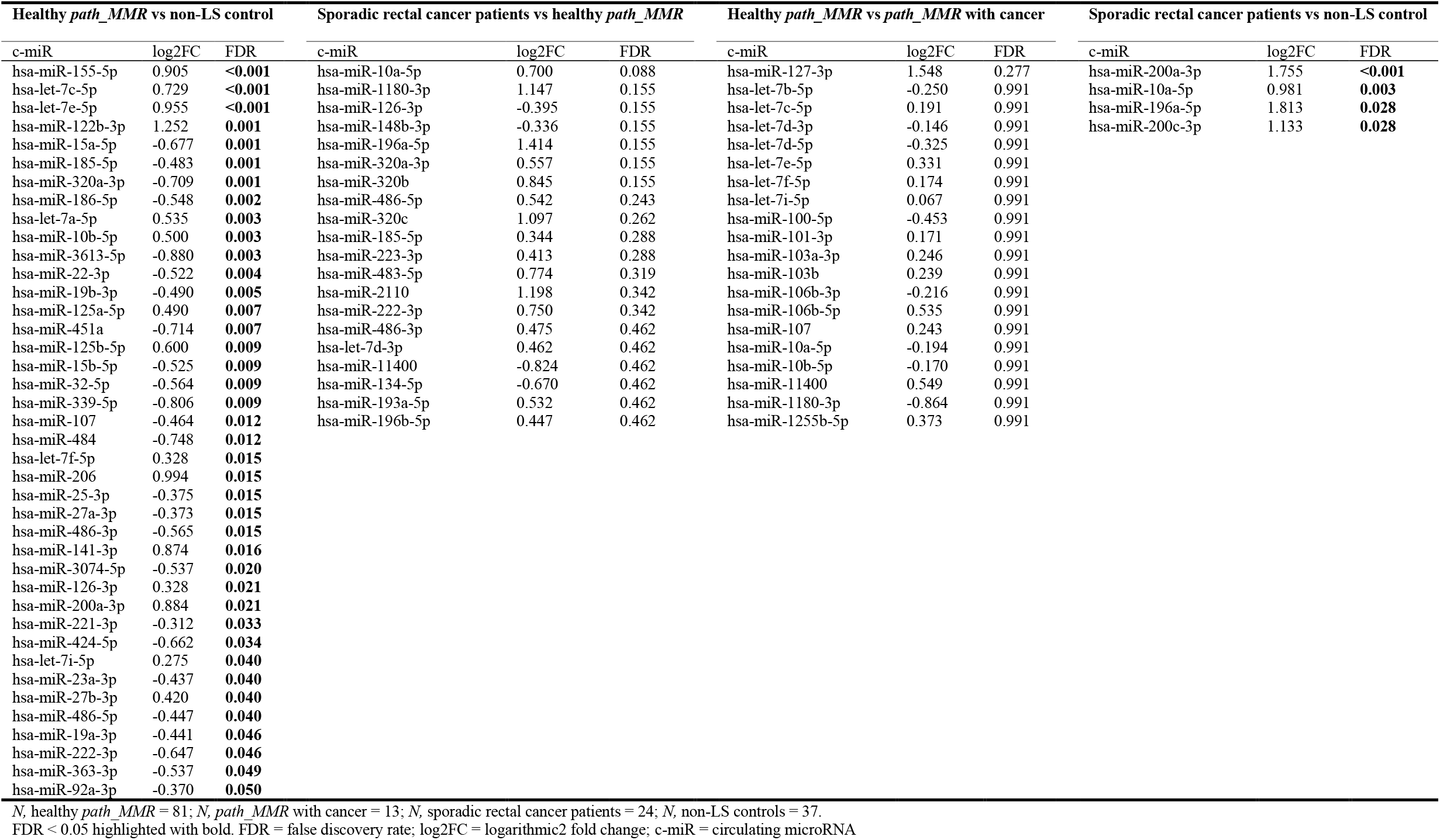
DE and non-DE c-miRs within and between the discovery and cancer cohorts.

**Figure 1.**
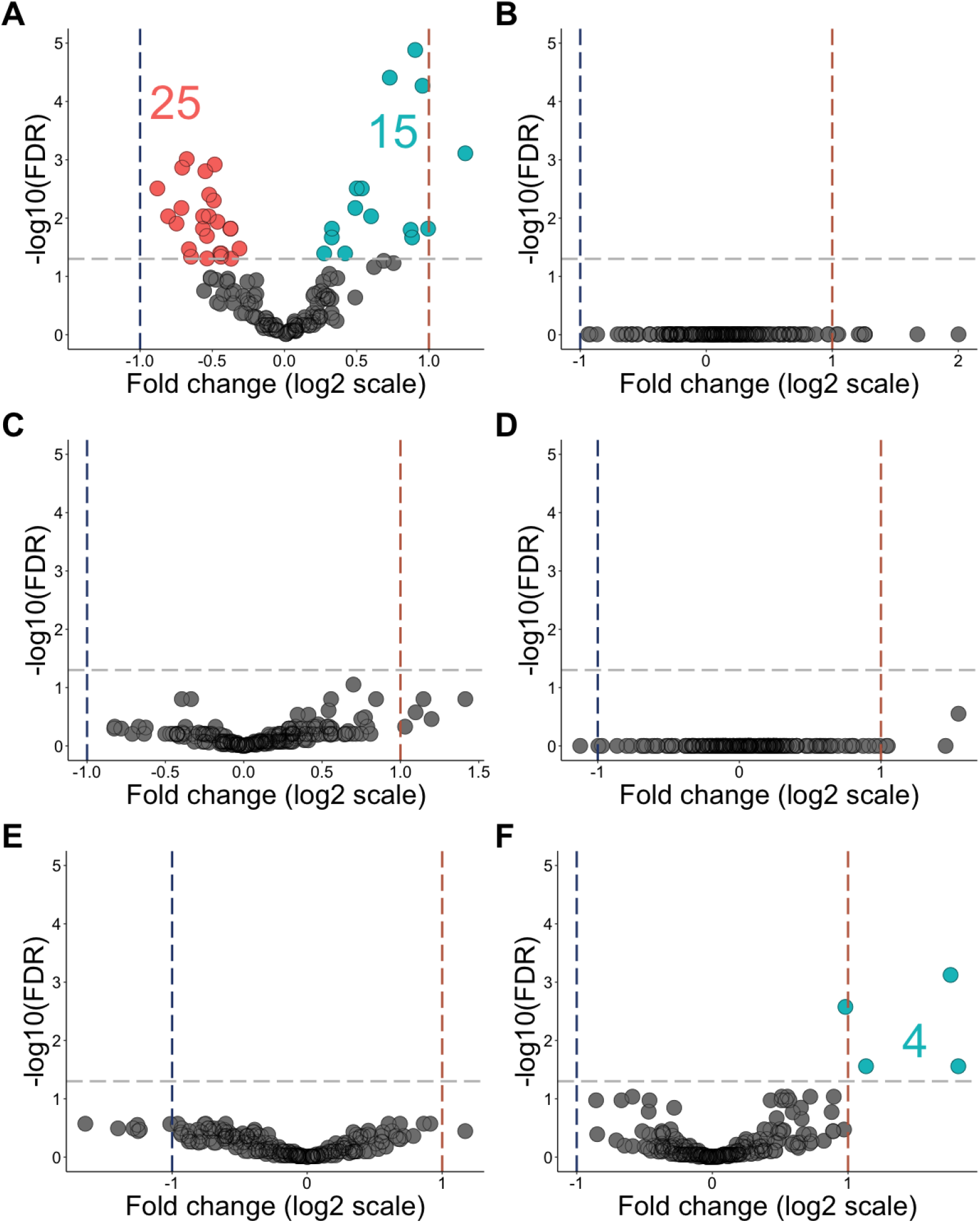
Healthy *path_MMR* carriers have a c-miRnome that differ from non-LS controls but resembles the c-miRnomes of patients with sporadic or hereditary cancer. **A**, DE c-miRs in healthy *path_MMR* carriers vs non-LS controls. **B**, DE c-miRs in sporadic rectal cancer patients vs *path_MMR* carriers with cancer **C**, DE c-miRs in healthy *path_MMR* carriers vs sporadic rectal cancer patients. **D**, DE c-miRs in healthy *path_MMR* carriers vs *path_MMR* carriers with cancer. **E**, DE c-miRs in *path_MMR* carriers with cancer vs non-LS controls. **F**, DE c-miRs in sporadic rectal cancers patients vs non-LS controls. Blue dash lines indicate negative fold change of expression, red dash line indicate positive fold change of expression and grey dash line indicate FDR <0.05. Downregulated c-miRs are highlighted in red, upregulated c-miRs are highlighted in cyan and non-significantly expressed c-miRs are highlighted in grey. Dots represents c-miRs. c-miR = circulating microRNA; FDR = false discovery rate.

To understand this phenomenon further, we explored whether the *path_MMR* carriers who currently have cancer also display unique c-miR expression. By using tumor samples, Balaguer et al. have shown that miR expression can distinguish LS tumors from sporadic tumors with microsatellite instability (14). To test if we can similarly reveal differences in c-miRs, we first inspected c-miRnomes within the cancer cohort but did not find any differences (Fig. 1B & Supplementary file 1, Table S2), thus suggesting a mutual c-miR response among the cancer types. Furthermore, our second analysis scheme comparing healthy *path_MMR* carriers to sporadic rectal cancer patients (Fig. 1C & Table 2), our third analysis scheme comparing healthy *path_MMR* carriers to *path_MMR* carriers with cancer (Fig. 1D & Table 2) and our fourth analysis scheme comparing *path_MMR* carriers with cancer to healthy non-LS controls (Fig. 1E & Supplementary file 1, Table S2) were also unable to detect DE c-miRs. These observations imply that c-miRnomes within our dataset can’t discern healthy *path_MMR* carriers from cancer patients with or without dMMR.

Several DE c-miRs have been implicated to sporadic cancer progression (41,42). To study this in our dataset, we compared sporadic rectal cancer patients to non-LS controls. We found that hsa-miR-200a-3p, -10a-5p, -196a-5p and -200c-3p were significantly upregulated in sporadic rectal cancer patients differentiating them from non-LS controls (Fig. 1F & Table 2). All of these c-miRs have earlier been shown to associate with colorectal cancer, and of them, hsa-miR-200a-3p was also significantly upregulated in healthy *path_MMR* carriers compared to non-LS controls with fold change of 0.88. In this analysis scheme, the fold change in hsa-miR-200a-3p was 1.76, indicating significantly higher expression compared to the healthy non-LS controls (Table 2).

Taken together, our findings imply that healthy *path_MMR* carriers have a systemic c-miRnome that separates them from healthy non-LS persons but resemble the c-miRnome of cancer patients with or without dMMR. Thus, these findings suggest that sporadic and dMMR-directed carcinogenesis share common miR-targeted biological pathways where potential alterations may produce a detectable c-miR signature in the healthy *path_MMR* carriers.

### Dimension reduction analysis of multiple traits was unable to discern *path_MMR* carriers from sporadic rectal cancer patients

We did not identify DE c-miRs between *path_MMR* carriers and sporadic rectal cancer patients. Therefore, by using the expression data of all 228 c-miRs shared between the discovery and cancer cohorts, we performed a dimension reduction analysis with t-SNE to identify possible subpopulations within *path_MMR* carriers and sporadic rectal cancer patients. First, we investigated if phenotypic traits such as being *path_MMR* carrier, current cancer status, cancer history or *path_MMR* variant type, would reveal clustering of samples, but did not find any clear patterns (Fig 2 A-D). We also investigated if age or BMI would be the discerning traits, but they also failed to reveal any clustering (Fig. 2E-G). Finally, sex and the sequencing batch did not form clusters within our dataset (Fig. 2H & I). Taken together, the t-SNE analysis supported the DE c-miR findings and was not able to differentiate *path_MMR* carriers from sporadic cancer patients, which may be an indicative of shared c-miR-mediated regulation as seen in the DE-analyses.

**Figure 2.**
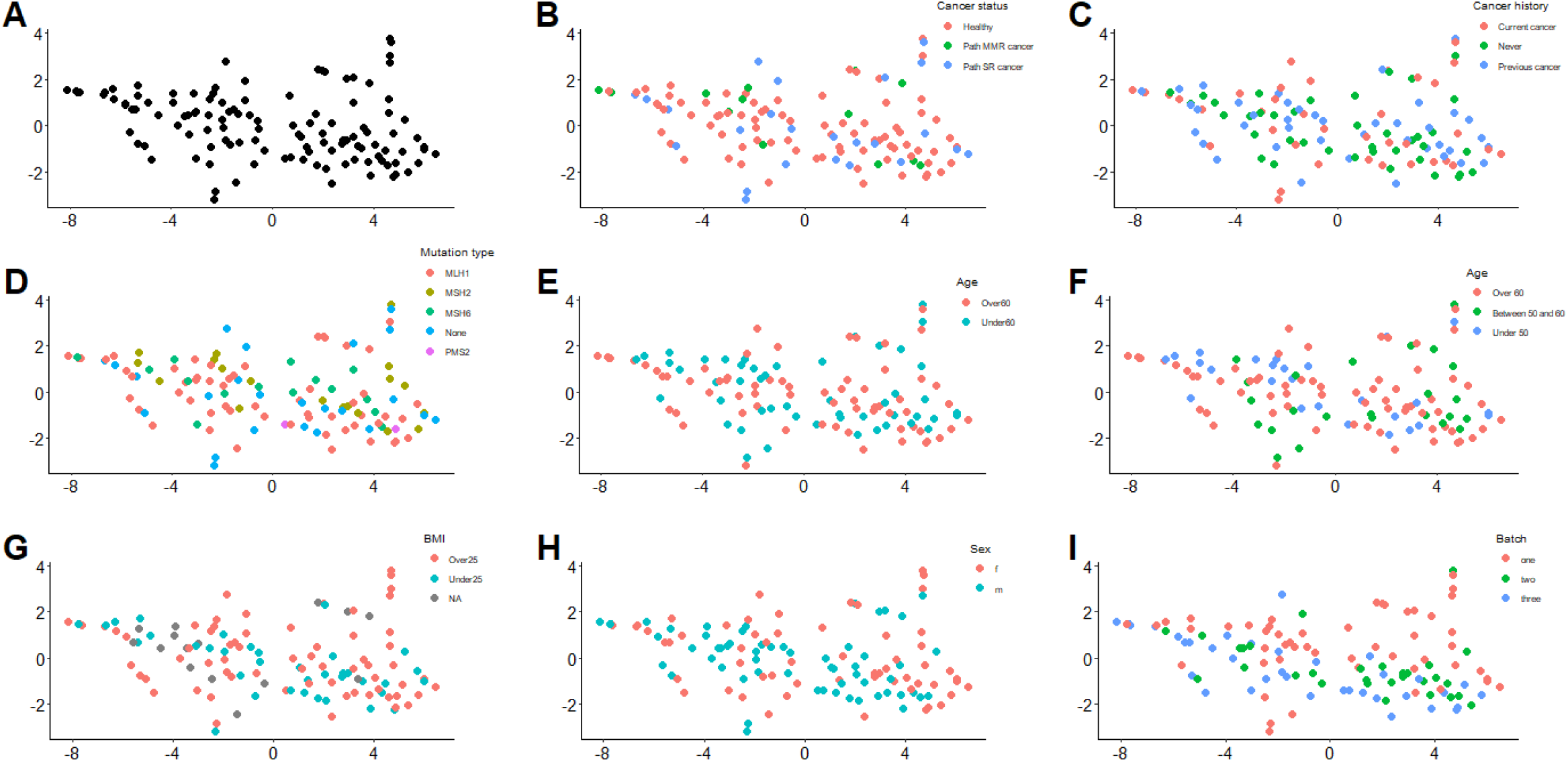
Dimension reduction analysis of multiple traits was unable to discern *path_MMR* carriers from sporadic rectal cancer patients. **A**, *Path_MMR* carriers and sporadic rectal cancer patients. **B**, Cancer status. Healthy = cancer-free path_MMR carriers; path_MMR cancer = path_MMR carriers with cancer; path SR cancer = sporadic rectal cancer patients. **C**, Cancer history. Current cancer = has cancer currently; Never = currently healthy, never had cancers; Previous cancer = currently healthy, had had cancer or multiple cancers; **D**, *path_MMR* variant. **E**, Dichotomous age. Over 60 = persons >60-years of age; Under 60 = persons <60-years of age. **F**, Non-dichotomous age. Over 60 = persons >60-years of age; Between 50 and 60 = persons between 50 to 60-years of age; Under 50 = persons <50-years of age **G**, BMI. Over 25 = persons with BMI>25; Under 25 = persons with BMI<25; NA = no reported BMI. **H**, Sex. M = males; F=females **I**, Batch effect of three separate sequencing runs in running order. All t-SNE plots are 2D constructions. Dots represent study subjects. BMI = body mass index.

### Pathway analysis revealed putative c-miR-target genes that are linked to biological processes and pathways associated with cancer

To further evaluate our hypothesis that healthy *path_MMR* carriers might have a c-miRnome that resembles the c-miRnome of cancer cohort due to shared miR-targeted biological pathways, we next investigated what are the target genes of the observed DE c-miRs. We also inspected what biological processes and pathways these target genes associate with. With mirWalk, we used random-forest-based approach to predict the target genes using databases with experimental validation and high confidence of reported miR-target gene interactions. MirWalk identified a total of 1,731 miR-target gene interactions with 508 distinct putative target genes for 32 out of 40 observed DE c-miRs from discovery cohort analysis (Supplementary file 4).

We then performed mirWalk-GSEA analysis on the 508 predicted target genes to explore what functional roles the DE c-miRs might possess. The GSEA analysis revealed 195 distinct significantly enriched biological processes (Supplementary file 5). To identify the key biological processes, we then narrowed the given output list based on FDR and the number of involved target genes to the top 30 most significantly enriched biological processes. Most of the discovered biological processes were linked to apoptosis, regulation of transcription, cell cycle, cell proliferation, DNA damage and signal transduction (Fig. 3A). We observed considerable overlap between the identified pathways since 208 out of 508 identified distinct c-miR-target genes contributed to these top biological processes (Supplementary file 5). *TGFBR1, CDKN1A, IGF1, TRAF6* and *BCL2* genes were present in most of the observed biological processes along with several other genes (Supplementary file 5). The performed *in silico* target analysis showed that *TGFBR1* was targeted by hsa-miR-27b-3p, *CDKN1A* and *IGF1* were targets of hsa-let-7e-5p, *TRAF6* was targeted by hsa-miR-125a-5p and *BCL2* was targeted by hsa-miR-125b-5p and hsa-miR-15b-5p (Supplementary file 4). Of these c-miRs, all except hsa-miR-15b-5p were upregulated in *path_MMR* compared to controls (Table 3).

**Table 3.**
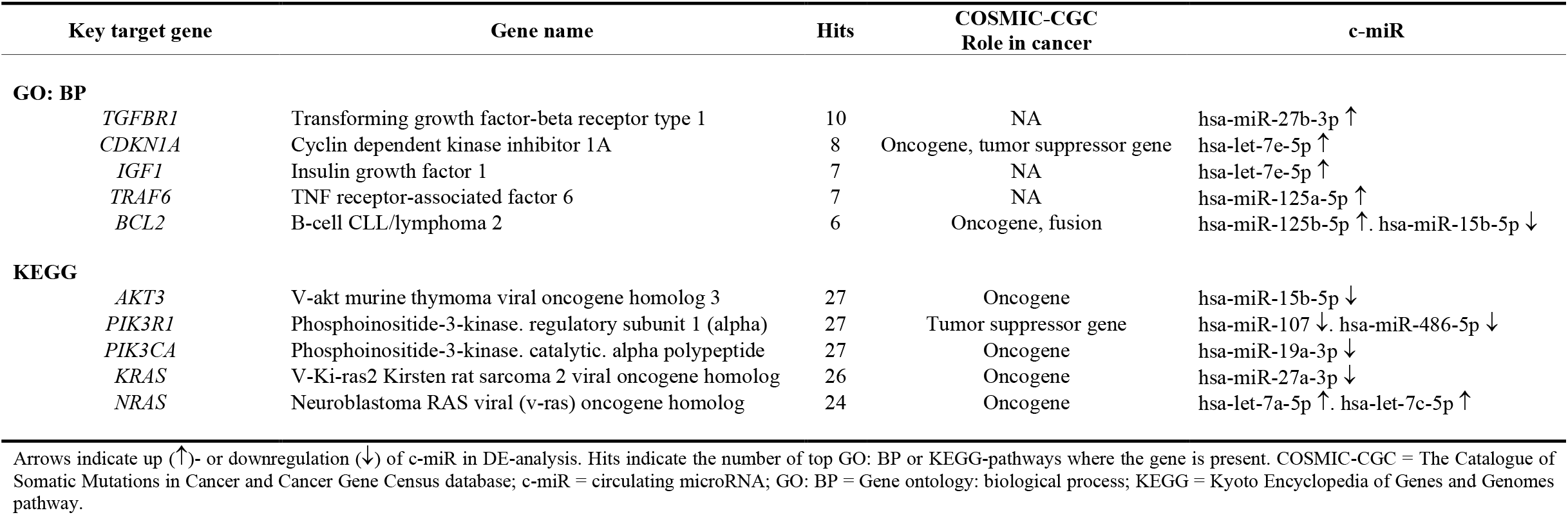
Key target genes of DE c-miRs in healthy *path_MMR* carriers compared to non-LS controls.

**Figure 3.**
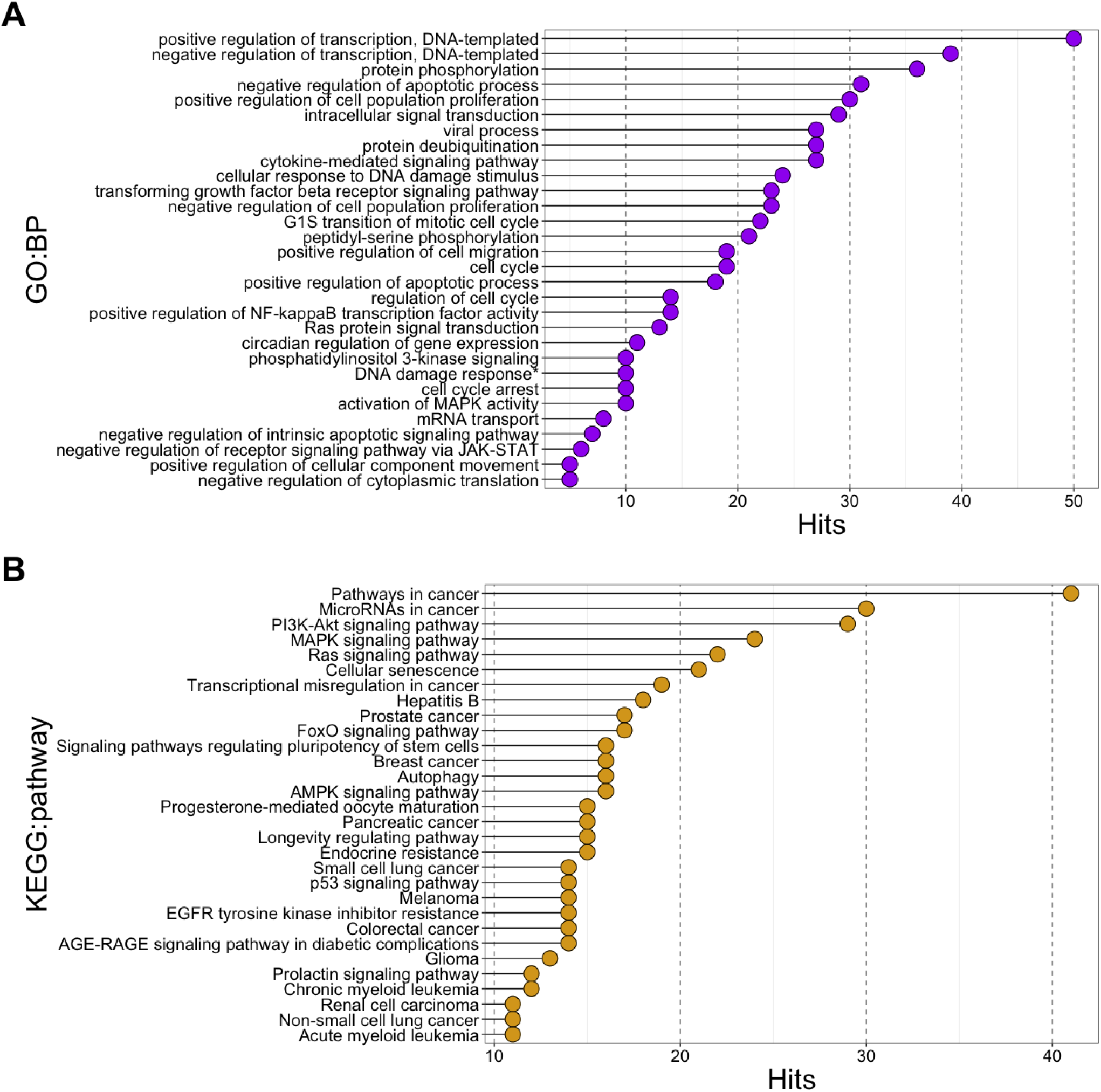
Pathway analysis revealed putative c-miR-target genes that are linked to biological processes and pathways associated with cancer. **A**, top 30 most enriched biological processes annotated to the identified target genes of 40 DE c-miRs found in healthy *path_MMR* carriers. FDR = false discovery rate; GO: BP = Gene Ontology: Biological process; Hits = number of target genes annotated to the biological process. *Signal transduction by p53 class mediator resulting in cell cycle arrest. **B**, top 30 most enriched KEGG pathways annotated to the identified target genes of 40 DE c-miRs found in healthy *path_MMR* carriers. c-miR = circulating microRNA; FDR = false discovery rate; KEGG = Kyoto Encyclopedia of genes and genomes pathway; Hits = number of target genes annotated to the pathway.

Next, we explored how the c-miR-target genes interact with KEGG pathways. GSEA analysis of the same gene set discovered 89 significantly enriched KEGG biological pathways (Supplementary file 6). Again, to focus on the possible key pathways, we narrowed the output list to the top 30 of the most significant pathways based on similar parameters than in the previous analysis. A great majority of the discovered pathways linked to cancer, cancer signaling and cellular aging (Fig. 3B). Of the 508 predicted target genes, 113 were involved in the discovered top KEGG pathways (Supplementary file 6). *AKT3, PIK3R1* and *PIK3CA* genes were involved in 27 out of 30 KEGG pathways, whereas *KRAS* had 26 and *NRAS* had 24 hits, respectively (Supplementary file 6). *AKT3* was targeted by hsa-miR-15b-5p, *PIK3R1* was targeted by hsa-miR-107 and hsa-miR-486-5p, *PIK3CA* was targeted by hsa-miR-19a-3p, *KRAS* was targeted by hsa-miR-27a-3p and *NRAS* was a target of hsa-let-7a and -7c-5p (Supplementary file 4). Of these c-miRs, all except hsa-let7a and -7c, were downregulated in *path_MMR* compared to controls (Table 3).

As these key target genes were interacting in the majority of the identified cancer-associated biological processes and pathways, we then explored and validated their potential carcinogenic roles. We submitted the gene set to COSMIC-CGC database and found that *BCL2, AKT3, PIK3CA, KRAS* and *NRAS* possess oncogenic functions, whereas *CDKN1A* is a potential oncogene or tumor suppressor gene and *PIK3R1* functions as a tumor suppressor gene (Table 3). All these genes have well-documented roles in multiple tumor types, including colorectal cancer, and with most having functions in hallmarks of cancer (43). Of the target gene set, *TGFBR1, IGF1* nor *TRAF6* were not included in COSMIC-CGC database. These results not only support our hypothesis that the observed resemblance of the c-miRnomes between *path_MMR* carriers and sporadic rectal cancer patients can be due to shared biological processes and pathways, but also support our GSEA findings by showing that the reciprocal factor behind resemblance is due to cancer progression governed by well-known oncogenes and tumor suppressor genes.

Taken together, our *in silico* analysis shows that the c-miRs in hsa-let-7 family, as well as hsa-miR-15b-5p, hsa-miR-19a-3p, hsa-miR-27a-3p and -27b-3p, hsa-miR-107, hsa-miR-125b-5p and hsa-miR-486-5p can be regulators of key target genes that are ubiquitous in cancer-associated biological processes and pathways. These findings imply that the altered c-miRnome expression pattern of cancer-free *path_MMR* carriers may hold predictive value by tracking potential oncogenic stress caused by dMMR-driven distortions.

## Discussion

Our study pioneered in characterizing the systemic c-miRnomes of *path_MMR* carriers. By utilizing high throughput sequencing, a total of 228 distinct c-miRs common to all study subjects were detected. Of these, we showed healthy *path_MMR* carriers to have an exclusive c-miRnome of 40 DE c-miRs that differs from non-LS-controls, but that does not differ from the c-miRnome of cancer patients with or without dMMR. Our c-miR expression analysis combined with *in silico* tools revealed that the observed resemblance in the c-miRnomes is possibly caused by distortions in several biological networks that are governed by well-known oncogenes and tumor suppressor genes, thus suggesting that c-miRnome could be used to track potential oncogenic stress at cancer-free state.

There is a growing interest in exploiting miRs as cancer biomarkers. Balaguer et al. studied miRs that were extracted from tumors of *path_MMR* carriers and sporadic colorectal cancer patients with verified microsatellite instability and normal tissue samples (14,18). They used a set of > 700 miR-probes with microarray analysis and detected hundreds of DE miRs among the tissue samples, showing that LS tumors can be separated from sporadic tumors with microsatellite instability, as well as that suspected LS samples discern from confirmed LS samples. Aligned with their study, we also showed that different *path_MMR* variants do not display unique c-miR expression thus implying a shared systemic response. However, we could not pinpoint DE c-miRs that would distinguish *path_MMR* carriers from sporadic cancer patients although we did, as well as in numerous other studies, detect a c-miR signature unique to sporadic cancer patients when compared to healthy non-LS controls. The observation that *path_MMR* carriers do not differ from sporadic cancer patients in their c-miRnome was also supported by our t-SNE analysis that didn’t reveal any clustering within our dataset based on several variables. The reason behind the substantial difference in DE c-miR numbers between our and the study by Balaguer et al. is likely explained by the study setting, used specimen type and methodology. In our study, the DE c-miRs were sequenced from the circulation of cancer-free persons where such a robust c-miR signature is not presumably detected when compared to miRs at the site of pathology.

Furthermore, Balaguer et al. detected several DE miRs with diagnostic potential in LS, including hsa-miR-125b-5p, -137, -622, -192 and -1238, whereas Zhou et al. displayed that hsa-miR-137, -520e and -590-3p are indicatives of LS by using a subset of *path_MMR* cancer tumor samples and normal tissue samples from the study by Balaguer et al. (17). We did not find significant overlapping of DE miR content between our c-miRs and tumor-miRs from those studies, except for hsa-miR-125b-5p, that was also identified by Balaguer et al. Aberrant expression of hsa-miR-125b-5p has been reported for multitude of cancer types and it has been implied to serve as a circulating cancer biomarker by targeting apoptosis-regulating oncogene *BCL2* (44).

The most significant DE c-miR in our setting was hsa-miR-155-5p, followed by hsa-let-7c-5p and -7e-5p, -122b-3p and 15a-5p, which all except hsa-miR-15a-5p were upregulated in healthy *path_MMR* carriers. Valeri et al. demonstrated that hsa-miR-155-5p targets several MMR-genes and further demonstrated that overexpression of hsa-miR155-5p downregulates *MLH1* and *MSH2* in colorectal cancer cell lines (15). Within this concept, our DE findings also support the role of hsa-miR-155p modulation in LS pathogenesis even though the performed *in silico* analysis could not identify MMR-genes as targets of hsa-miR-155-5p. miRs in hsa-let-7 family have been suggested to increase colorectal cancer risk in *path_MMR* carriers with proficient MMR by lowering the expression of *TGFBR1* haplotype (45). We found hsa-let-7 family to target *TGFBR3* and hsa-miR-27b-3pto target *TGFBR1*. We did not find target genes for hsa-miR-122b-3p, and thus role of this c-miR in our study remained unknown. Previous studies have linked hsa-miR-15a-5p to sporadic endometrial cancer (46) and colorectal cancer (47), both being hallmark cancers of LS. In our analysis, hsa-miR-15a-5p was seen to target several genes, including known oncogenes and tumor suppressor genes such as *CCND1, CDK6* and *DICER1*, thus suggesting biomarker potential also in LS.

miRs have critical functions across various biological processes and pathways involved in carcinogenesis. We found 508 putative target genes for 32 out of 40 observed DE c-miRs that associate with several pathways common to cancer. In addition to above mentioned c-miRs, we also identified several other c-miRs that could be key regulators in dMMR-driven carcinogenesis. All these c-miRs were observed to target several well-known oncogenes and tumor suppressor genes such as *KRAS, NRAS, PIK3RI*, and *PIK3CA*, that were significantly enriched in our pathway analysis. Supported by our DE-analysis, the observation that these identified DE c-miRs target known oncogenes and tumor suppressors, could indicate upregulation of the oncogenes and consequently downregulation of the tumor suppressor genes. However, since we studied cell-free c-miRs without possibility to investigate expression levels of their putative target genes, this suggestion remains hypothetical. Unfortunately, c-miRs are not easily tracked where tracking of c-miRs would provide us clues to what tissues they will be affecting and where to seek further signs for cancer development. Matching pairwise tissue samples to observed c-miRs could help elucidate these issues but we had no possibility to do so. Nevertheless, our exploratory findings indicate that *path_MMR* carriers display oncogenic stress even when they are cancer-free, but more studies are needed to verify our results and to show if they have true power as a biomarkers of early cancer development.

A major strength of our study is that the study subjects had undergone comprehensive screenings of LS-predisposing mutations, with ascertainment utilizing Amsterdam and Bethesda clinical criteria and cascade testing. Also, instead of *a priori* chosen gene panel, we conducted a systemic level investigation of c-miRnome, which provides a more comprehensive view of how already identified c-miRs and putative target genes contribute to distorted biological networks in sporadic and hereditary cancer. For example, our findings allow construction of c-miRnome-target gene collection to be explored for potentially distorted biological networks associated with dMMR. Also, it can be used for establishing candidate hypotheses to drive further research and for further exploratory c-miR analyses of potential contributing gene clusters not previously discovered. Finally, the bioinformatic analyses in our study were performed in precise detail according to the latest knowledge using state-of-the-art tools and algorithms.

Our study has potential pitfalls. Although largest to date, the study sample was relatively small especially in the cancer cohort, which could have reduced the statistical power of DE-analyses. Regarding the methodology, there is no conclusive rule which sequencing depth should be aimed at when assessing DE of c-miRs. In our study, the aimed mean sequencing depth was 5M reads per sample, but the achieved mean sequencing depth was 3.2M reads due to under-clustering issues in sequencing. The under-clustering might have affected c-miR detection by favoring highly expressed c-miRs and thus resulting in over-presentation of these c-miRs and under-presentation or masking of c-miRs with low expression and potential cancer- or dMMR-relevant functions. A common issue with c-miRs is the identification of their primary and target locations, and alike in many other studies, we did not track the observed c-miRs to certain locations, which introduce a certain degree of uncertainty over the interpretations of the observations. Unfortunately, our efforts to validate DE findings with RT-qPCR were not completely successful when using an independent validation cohort, although we observed a trend of parallel expression in both cohorts in eight out of nine validation c-miRs. Overall considerable variation in c-miR expression levels were detected with both methods and cohorts, that could explain why significant differences between groups in the smaller validation cohort were not detected.

Our exploratory study produced novel insight by elucidating the potential roles of c-miR-target gene interactions in LS, thus paving way for the future investigation of c-miRs in monitoring the risk stratification patterns during the risk-reducing clinical surveillance and possible cancer management. A future goal is to determine whether the longitudinal change or development of c-miRnomes appears in conjunction with cancer incidence and treatment. The biological basis for aberrant c-miR expression between *path_MMR* carriers and non-LS controls remains a clinical question to be elucidated also in the future work.

## Authors’ contributions

**T. Sievänen:** Formal analysis, investigation, methodology, software, validation, visualization, writing original draft, -review and -editing. **T**.**-M. Korhonen:** Formal analysis, methodology, software, visualization, writing-review and -editing. **T. Jokela:** Methodology, software, validation, visualization. **M. Ahtiainen:** Methodology, writing-review and -editing. **L. Lahtinen:** Methodology, writing-review and -editing. **T. Kuopio:** Methodology, writing-review and -editing. **A. Lepistö:** Writing-review and - editing. **E. Sillanpää:** Conceptualization, supervision, writing-review and -editing. **J-P. Mecklin:** Conceptualization, resources, writing-review and -editing. **T**.**T. Seppälä:** Conceptualization, funding acquisition, resources, supervision, writing-review and -editing. **E**.**K. Laakkonen:** Conceptualization, data curation, funding acquisition, project administration, resources, supervision, writing-review and - editing.

## Acknowledgements

We acknowledge the support from Biosciences team of IT Center for Science Finland (CSC) for providing HPC-resources for our data analytics.

## Author’s Disclosures

The authors declare no potential conflicts of interest.

## Grants

E.K. Laakkonen was supported by the Päivikki and Sakari Sohlberg Foundation. E. Sillanpää was supported by the Academy of Finland research fellowship (grant number: 341750). T.T. Seppälä was supported by Finnish Medical Foundation, Emil Aaltonen Foundation, Jane and Aatos Erkko Foundation, Sigrid Juselius Foundation, Finnish Cancer Foundation, Relander Foundation, Academy of Finland (grant number: 338657, HUS State Research Funds (TYH2021123 and TYH2022323) and iCAN Digital Precision Medicine Flagship. J-P. Mecklin was supported by Jane and Aatos Erkko Foundation, Finnish Cancer Foundation and KYS State Research Funds.

## Notes

### Competing Interest Statement

The authors have declared no competing interest.

