## Supplementary file 1 for "Systemic circulating microRNA landscape in Lynch syndrome"

This supplementary file contains 1) small-RNA sequencing summary statistics, 2) microRNA discovery summary, 3) differential expression analysis results and 4) RT-qPCR validation summary.

### 1. Small-RNA sequencing summary statistics

The summary statistics of all sequencing runs are listed in Table 1. Samples that had < 1M raw reads were excluded from the downstream analysis (n=5). A total of 155 samples were sequenced successfully in three separate sequencing runs with minor under-clustering affecting the runs. The mean raw read count of the experiment was 3,761,804 M reads per sample and after adapter removal, trimming and filtering of low-quality data the mean clean read count was 1,601,265 M reads per sample. On average, 1,150,017 clean reads per sample mapped to human microRNAs (miRs) resulting in mean alignment rate of 63 %.

**Table S1**. Sequencing summary statistics of all sequencing runs.

| **SEQUENCING RUN I** | | | | |
| --- | --- | --- | --- | --- |
| **Parameter** | **Lynch syndrome** | **Sporadic rectal cancer patient** | **Control** | **Total** |
| N | 41 | 10 | 10 | 61 |
| Raw read count (mean[count], ± SD) | 2,997,064  (677,455) | 3,341,588 (1,586,332) | 3,508,312  (1,180,659) | 3,282,321  (455,310) |
| Clean read count (mean[count], ± SD) | 1,760,483  (518,578) | 1,123,486  (542,040) | 1,814,271  (820,051) | 1,566,080  (167,693) |
| MicroRNA-aligned read count (mean[count], ± SD) | 1,009,995  (447,449) | 678,927  (324,164) | 1,079,379  (592,904) | 922,767  (134,522) |
| Alignment-% (mean [%]) | 55.5 % | 61.0 % | 57.5 % | 58.0 % |
| **SEQUENCING RUN II** | | | | |
| **Parameter** | **Lynch syndrome** | **Sporadic rectal cancer patient** | **Control** | **Total** |
| N | 24 | 6 | 7 | 37 |
| Raw read count (mean[count], ± SD) | 3,850,746  (1,514,056) | 4,593,992  (743,092) | 3,796,429  (1,298,281) | 4,080,389  (397,739) |
| Clean read count (mean[count], ± SD) | 1,444,305  (950,828) | 1,090,272  (624,565) | 1,732,170  (924,363) | 1,422,249  (181,212) |
| MicroRNA-aligned read count (mean[count], ± SD) | 983,377  (686,158) | 669,817  (401,775) | 1,113,962  (688,652) | 922,385  (164,913) |
| Alignment-% (mean [%]) | 65.8 % | 58.9 % | 63.2 % | 62.6 % |
| **SEQUENCING RUN III** | | | | |
| **Parameter** | **Lynch syndrome** | **Sporadic rectal cancer patient** | **Control** | **Total** |
| N | 29 | 8 | 20 | 57 |
| Raw read count (mean[count], ± SD) | 3,884,176  (1,663,080) | 4,605,843  (1,008,139) | 4,782,816  (1,419,636) | 4,424,278  (331,044) |
| Clean read count (mean[count], ± SD) | 1,675,657  (1,190,841) | 1,256,684  (554,348) | 1,733,051  (1,134,394) | 1,555,131  (352,317) |
| MicroRNA-aligned read count (mean[count], ± SD) | 1,219,297  (1,005,301) | 708,023  (388,872) | 1,256,710  (862,318) | 1,061,344  (322,640) |
| Alignment-% (mean [%]) | 70.5% | 57.0% | 71.2% | 66.2 % |
| **SEQUENCING RUN I-III** | | | | |
| **Parameter** | **Lynch syndrome** | **Sporadic rectal cancer patient** | **Control** | **Total** |
| N | 94 | 24 | 37 | 155 |
| Raw read count (mean[count], ± SD) | 3,577,329  (531,282) | 4,180,474  (431,202) | 4,029,185  (119,493) | 3,761,804  (1,395,080) |
| Clean read count (mean[count], ± SD) | 1,626,815  (340,682) | 1,156,814  (44,520) | 1,759,831  (160,107) | 1,601,265  (886,932) |
| MicroRNA-aligned read count (mean[count], ± SD) | 1,070,890  (279,891) | 685,589  (41,587) | 1,150,017  (136,572) | 1,035,926  (698,458) |
| Alignment-% (mean [%]) | 63.9 % | 59.0 % | 64.0 % | 63.0 % |

### 2. MicroRNA discovery summary

The alignment produced 1349 distinct c-miRs within the discovery cohort (supplementary file 3). After filtering out the genes with low expression (<1 count per million in 70% of samples per treatment group), 228 c-miRs were left for differential expression analysis (supplementary file 3). All the 228 c-miRs were observed in all three treatment groups (Figure 1). Top 5 c-miRs with the highest raw counts (highest sequencing depth) as a proportion of all c-miR counts per sample for Lynch syndrome (LS), sporadic rectal cancer patients (SRME) and control (CTRL) groups are presented in Figures 2, 3 and 4, respectively. Hsa-miR-16-5p had the highest raw count in all groups followed by hsa-let-7a-5p, hsa-let-7b-5p, hsa-miR-122-5p (SRME only) and hsa-miR-223-3p and hsa-miR-451a.

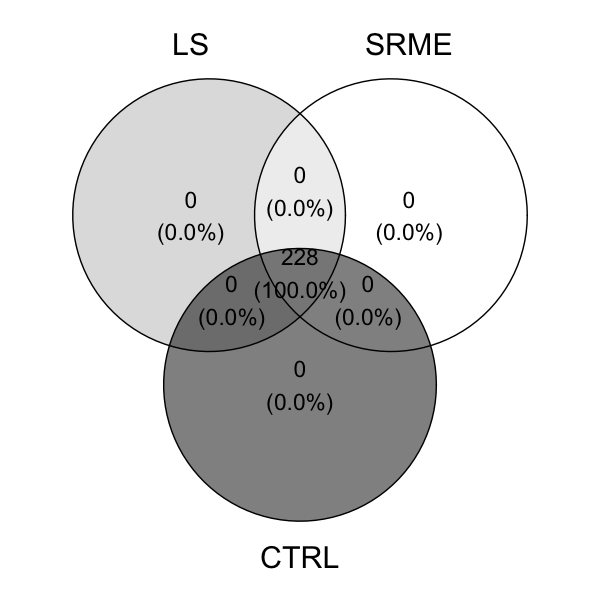

**Figure S1.** Venn diagram of c-miR distribution among treatment groups. LS = Lynch syndrome, SRME = sporadic rectal cancer patients, CTRL = control.

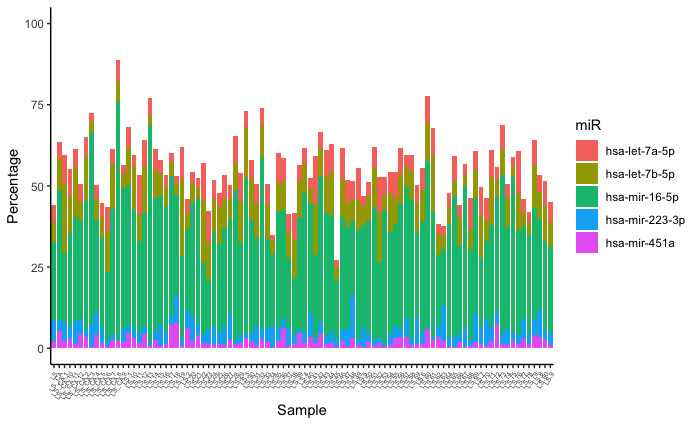

**Figure S2.** Top 5 c-miRs with the highest raw counts (highest sequencing depth) as a proportion of all c-miR counts per sample in Lynch syndrome group. Samples are shown on the x-axis and percentages on the y-axis.

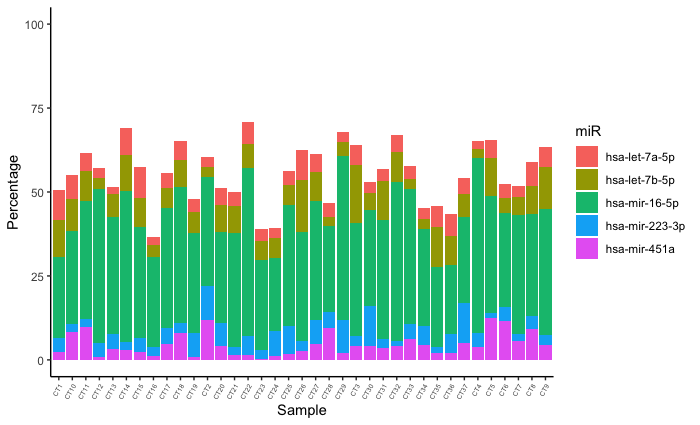

**Figure S3.** Top 5 c-miRs with the highest raw counts (highest sequencing depth) as a proportion of all c-miR counts per sample in control group. Samples are shown on the x-axis and percentages on the y-axis.

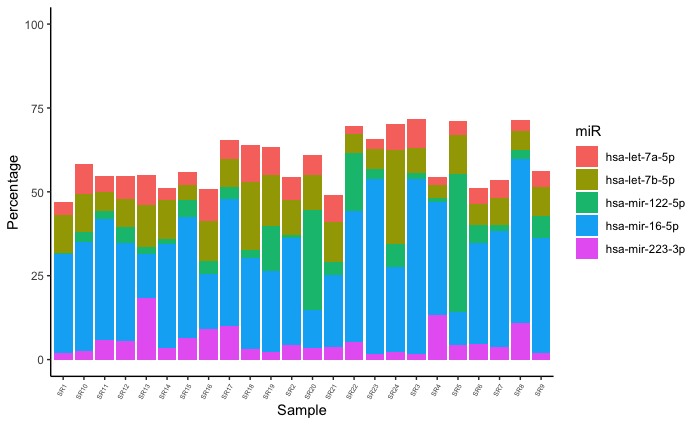

**Figure S4.** Top 5 c-miRs with the highest raw counts (highest sequencing depth) as a proportion of all c-miR counts per sample in sporadic rectal cancer patient group. Samples are shown on the x-axis and percentages on the y-axis.

3. **Differential expression analysis**

**Table S2**. Non-differentially expressed c-miRs within and between the discovery and cancer cohorts.

| **Sporadic rectal cancer patients vs *path_MMR with cancer*** | | |  | ***Path_MMR* with cancer vs non-LS control** | | |
| --- | --- | --- | --- | --- | --- | --- |
| miR | log2FC | FDR |  | miR | log2FC | FDR |
| hsa-let-7a-5p | 0.009 | 0.984 |  | hsa-mir-10b-5p | 0.688 | 0.268 |
| hsa-let-7b-5p | 0.065 | 0.984 |  | hsa-mir-125a-5p | 0.603 | 0.268 |
| hsa-let-7c-5p | 0.223 | 0.984 |  | hsa-mir-127-3p | -1.643 | 0.268 |
| hsa-let-7d-3p | 0.295 | 0.984 |  | hsa-mir-144-3p | -0.959 | 0.268 |
| hsa-let-7d-5p | -0.229 | 0.984 |  | hsa-mir-16-5p | 0.863 | 0.268 |
| hsa-let-7e-5p | 0.261 | 0.984 |  | hsa-mir-32-5p | -0.749 | 0.268 |
| hsa-let-7f-5p | 0.237 | 0.984 |  | hsa-mir-361-3p | 0.913 | 0.268 |
| hsa-let-7g-5p | 0.195 | 0.984 |  | hsa-mir-423-3p | -0.603 | 0.268 |
| hsa-let-7i-5p | -0.109 | 0.984 |  | hsa-mir-15b-5p | -0.760 | 0.271 |
| hsa-mir-100-5p | -0.585 | 0.984 |  | hsa-mir-374a-5p | -1.014 | 0.271 |
| hsa-mir-101-3p | -0.255 | 0.984 |  | hsa-mir-181d-5p | -1.257 | 0.280 |
| hsa-mir-103a-3p | 0.136 | 0.984 |  | hsa-mir-15a-5p | -0.736 | 0.322 |
| hsa-mir-103b | 0.107 | 0.984 |  | hsa-mir-190a-5p | -0.695 | 0.322 |
| hsa-mir-106b-3p | -0.593 | 0.984 |  | hsa-mir-320a-3p | -0.649 | 0.322 |
| hsa-mir-106b-5p | 0.460 | 0.984 |  | hsa-mir-370-3p | -1.287 | 0.322 |
| hsa-mir-107 | 0.262 | 0.984 |  | hsa-mir-432-5p | -0.930 | 0.322 |
| hsa-mir-10a-5p | 0.284 | 0.984 |  | hsa-mir-451a | -0.730 | 0.322 |
| hsa-mir-10b-5p | 0.143 | 0.984 |  | hsa-mir-654-3p | -1.401 | 0.322 |
| hsa-mir-11400 | -0.085 | 0.984 |  | hsa-mir-107 | -0.615 | 0.339 |
| hsa-mir-1180-3p | 0.076 | 0.984 |  | hsa-mir-221-3p | -0.389 | 0.339 |

5. **RT-qPCR validation summary**

### RT-qPCR validation of the selected differentially expressed c-miRs was performed using independent validation cohort that is independent from discovery and cancer cohorts

Validation cohort (n=29) comprised of 14 healthy *path_MMR* carriers and 15 non-LS controls (Table S3). DE c-miRs identified in the discovery cohort with log2 fold change being significantly above or below the average and with mean count >100 counts were chosen to validate the sequencing results using an independent validation cohort. These inclusion criteria were chosen to increase the probability of detecting c-miRs with RT-qPCR which is less sensitive and more prone to noise than sequencing. After small-RNA isolation, cDNA was synthetized with miRCURY®LNA® RT kit (339340, Qiagen). cDNA synthesis was done from 12 µl of non-diluted template RNA and PCR protocol was carried out in standard thermocycler (Eppendorf). Transcripts levels were measured by using miRCURY LNA™ miRNA PCR assays (has-let-7e-5p, hsa-miR-141-3p, hsa-miR-155-5p, hsa-miR-206, hsa-miR-320a, hsa-miR-339-5p, hsa-miR-451a, hsa-miR-484, hsa-miR-3613-5p, 339350, Qiagen) and miRCURY LNA SYBR® Green kit (339346, Qiagen). One μl of 1:2 diluted cDNA was used per well and samples were run as triplicates. qPCR protocol was the following: 95°C (2 min, activation), 95°C (10 s), 56°C (60 s) with 40 cycles (CFX384™ Real-Time PCR Detection System, Bio-Rad). Fold expression was calculated using the formula 2(-ΔΔCt), where ΔCt(sample)- ΔCt (mean Ct from all samples), ΔCt is Ct (c-miR of interest)-Ct (mean Ct from the group) and Ct is the cycle at which the detection threshold is crossed. Samples with Ct values >35 were excluded from the analysis.

**Table S3.** Descriptive characteristics of study subjects in the validation cohort.

|  |  | **Validation cohort** | |
| --- | --- | --- | --- |
| **Variable** |  | ***Path_MMR,* healthy** | **non-LS, healthy** |
| **N** |  | 14 | 15 |
| **Sex** (N (%)) |  |  |  |
| Male |  | 6 (42.9) | 8 (53.3) |
| Female |  | 8 (57.1) | 7 (46.7) |
| **Age,** years (mean± SD) |  | 59.7 (11.9) | 57.3 (15.1) |
| **Body mass index,** kg/m^2^ (mean± SD)^#^ |  | 28.6 (7.9) | 30.7 (5.1) |
| ***Path_MMR*** (N (%)) |  |  |  |
| *MLH1* |  | 14 (100.0) | - |
| *MSH2* |  | 0 (0.0) | - |
| *MSH6* |  | 0 (0.0) | - |
| *PMS2* |  | 0 (0.0) | - |
| **Previous cancers** (N (%)) |  |  |  |
| Yes |  | 9 (64.2) | - |
| No |  | 5 (35.8) | - |

Gene expression of the selected c-miRs (hsa-let-7e-5p, hsa-miR-141-3p, hsa-miR-155-5p, hsa-miR-206, hsa-miR-320a, hsa-miR-339-5p, hsa-miR-451a, hsa-miR-484 and hsa-miR-3613-5p) did not display statistically significant differences between the discovery and validation cohorts (Fig. S5A). Pearson correlations of the c-miR expression fold changes between the discovery and validation cohorts showed that seven out of nine RT-qPCR validated c-miRs had corresponding fold changes in both cohorts (p<0.001, r = 0.985). Two c-miRs, hsa-miR-206 and -3613-5p, showed deviations (Fig. S5B). Hsa-miR-206 had positive fold change in both cohorts but the magnitude of the fold change value was substantially higher in the validation cohort compared to the discovery cohort. The fold change of hsa-miR-3613-5p differentiated between the cohorts by showing downregulation in discovery cohort but upregulation in validation cohort (Fig. S3B). Taken together, c-miR expression fold changes followed an analogous trend in both cohorts in 8 out of 9 validation c-miRs. The substantial variation in the overall c-miR expression within the validation cohort due to small sample size could have affected the statistical power and thus failed to verify our findings.

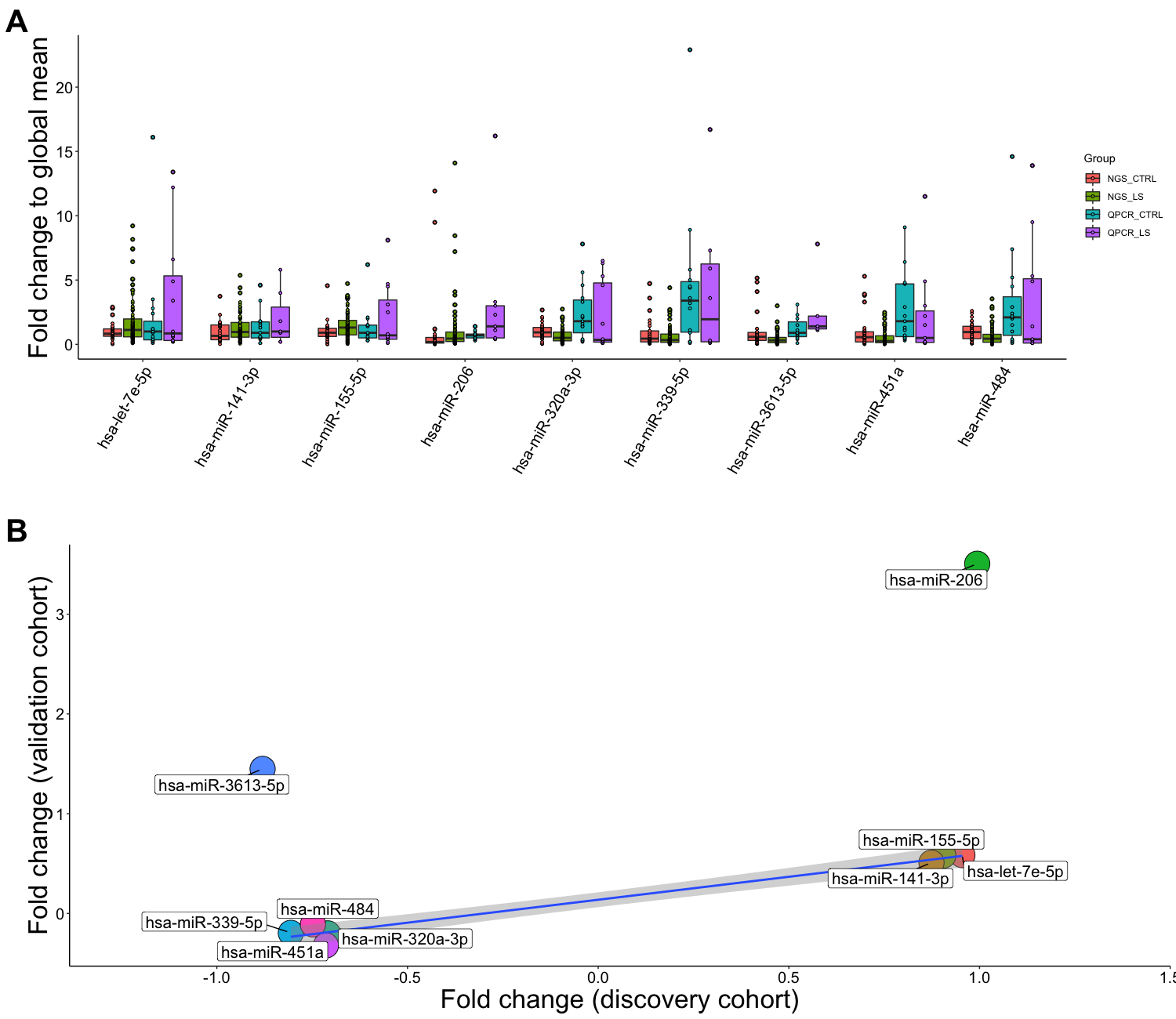

**Figure S5.** RT-qPCR validation revealed significant variation in c-miR expression in the validation cohort. **A,** distributions of expression fold changes of the nine selected c-miRs compared to global mean expression between the discovery and validation cohorts. Error bars represent SEM. Red = non-LS control samples from discovery cohort; Green = healthy *path_MMR* carriers from discovery cohort; Cyan = non-LS control samples from validation cohort; Purple = *path_MMR* carriers from validation cohort. **B,** Pearson correlations of expression fold changes of selected c-miRs between the discovery and validation cohorts. Blue line indicates the linear model fit of the 7 c-miRs which correlated between the experiments and grey cloud represents variation. c-miR = circulating microRNA.
