## Supplementary file 3 for "Systemic circulating microRNA landscape in Lynch syndrome": Supplementary_file_3.html

manuscriptAnalysis\_groomed.knit


Code 

- Show All Code
- Hide All Code

### **Code supplementary file**

###### Tero Sievänen & Tia-Marje Korhonen

#### *2022-02-03*

### 1 Introduction

This code supplementary file was used to perform all differential expression analyses between and within the discovery and cancer cohorts.

---

### 2 R-packages used

A variety of R packages was used for this analysis. All packages used are available from the Comprehensive R Archive Network (CRAN), Bioconductor.org, or Github.

---

### 3 Contact information

All the scripts have been written by Tero Sievänen (Uni. of Jyväskylä) and Tia-Marje Korhonen (Uni. of Jyväskylä).

---

### 4 Preprocessing and read mapping

#### 4.1 Preprocessing

FastQC was used for sequence quality control throughout the pipeline. Sequencing adapter removal, trimming and filtering was done with FASTX-Toolkit.

#### 4.2 Read mapping

Bowtie aligner was used for mapping the preprocessed high-quality reads to human miR-genome derived from miRBase v.22. A cut-off of >=1M reads per sample was applied before bioinformatic analysis. Mean raw read count was 3,761,804 M and 1,082,304 M was the mean c-miR read count across all samples. The mean alignment rate throughout the study population was 63%.

---

### 5 Analysis

#### 5.1 Step 1 - C-miR count data filtering

This script is used to make a filtered c-miR raw count file. Genes with low counts must be removed before DE-analysis, since their biological significance is low or might be false positives.

```
# Load packages
library(edgeR) # Package for differential gene expression analysis of RNA-seq data

# Import raw counts file
counts <- read.csv("rawCounts.tsv",header=TRUE,sep="\t")

# Import study design file
targets<- read.csv("phenodata.txt",sep="\t",header=TRUE)

# Add column names
colnames(counts) <-targets$Filename

# Filter phenodata based on sample type
Group1  <- which(targets$Type=="LS")
Group2  <- which(targets$Type=="CTRL")
Group3  <- which(targets$Type=="SRME")
ls <- targets[Group1,]
ctrl <- targets[Group2,]
srme <- targets[Group3,]

# Create a digital gene expression list (DGEList) using edgeR
myDGEList <- DGEList(counts = counts)

# Convert DGEList to counts per million (cpm)
cpm <- cpm(myDGEList)

# Keep only the c-miRs with >1 CPM in at least 70% of samples in a subgroup
keepers <- rowSums(cpm>1)>=108  #70% of 155

# Filter the subgroups (LS,CTRL and SRME)
cpm1 <- cpm[,which(colnames(cpm)%in%ls$Sample)]
cpm2 <- cpm[,which(colnames(cpm)%in%ctrl$Sample)]
cpm3 <- cpm[,which(colnames(cpm)%in%srme$Sample)]

keep1 <- rowSums(cpm1>=1) >= 65   #  70% of 94
keep2 <- rowSums(cpm2>=1) >= 25   #  70% of 37
keep3<- rowSums(cpm3>=1) >= 16   #  70% of 24

# Create a new filtered DGEList
myDGEList.filtered <- myDGEList[(keepers|keep1|keep2|keep3),]

# Create a new filtered raw counts matrix
FilteredCounts <- counts[which(rownames(counts)%in%rownames(myDGEList.filtered$counts)),]

# Create a new .txt file of filtered raw miR counts
write.table(FilteredCounts,"FilteredCounts.txt",sep="\t")
print("Step 1 completed!")
```

```
## [1] "Step 1 completed!"
```

---

#### 5.2 Step 2 - Sex difference within *path\_MMR* carriers, sporadic rectal cancer patients, and non-LS control group

This script is used to perform DE-analysis between sexes among *path\_MMR* carriers, sporadic rectal cancer patients (SRME) and non-LS control group (CTRL).

```
# Load packages
library(tidyverse) # Tidyverse is an opinionated collection of R packages designed for data science
library(gt) # Package for making static data tables
library(DESeq2) # Package for differential gene expression analysis of RNA-seq data
library(plotly) # Package for making interactive plots
library(DT) # Package for making interactive tables

# Read in the filtered c-miR counts table from step 1
counts <- read.csv("FilteredCounts.txt",header=TRUE,sep="\t")

# Add sample names (column names) to raw counts table
colnames(counts) <-targets$Filename

# Choose only the healthy path_MMR carriers (!= cancer | future_ca), n = 81
select <- which(targets$Type=="LS" & targets$Healthy_now=="YES")

# AND/ OR 
#select <- which(targets$Type=="SRME") # No sex difference
#select <- which(targets$Type=="CTRL") # No sex difference

# Create a new filtered counts file 
Counts <- counts[,select]

# New phenofile with only the variables of interest
Targets <- targets[select,]

# Setup design matrix for DE-analysis
condition <- as.character(Targets$Sex) # Condition of interest
batch <- as.character(Targets$NGS) # Batch effect
group_levels <- levels(as.factor(condition))
design <- data.frame(condition=as.factor(condition), batch=batch) # DE-design for the analysis
rownames(design) <- colnames(Counts)
dds <- DESeqDataSetFromMatrix(countData=Counts, colData=design, design = ~ batch + condition)

# DESeq2 DE-analysis of the condition of interest, batch effect taken into account
dds <- DESeq(dds)

# Display results
res <- results(dds, alpha=0.05) # Statistical significance at the level p=< 0.05
resOrdered <- res[order(res$padj),] # Order results based on adjusted p-value
summary(res)
```

```
## 
## out of 228 with nonzero total read count
## adjusted p-value < 0.05
## LFC > 0 (up)       : 0, 0%
## LFC < 0 (down)     : 2, 0.88%
## outliers [1]       : 0, 0%
## low counts [2]     : 0, 0%
## (mean count < 8)
## [1] see 'cooksCutoff' argument of ?results
## [2] see 'independentFiltering' argument of ?results
```

```
resOrdered
```

```
## log2 fold change (MLE): condition m vs f 
## Wald test p-value: condition m vs f 
## DataFrame with 228 rows and 6 columns
##                         baseMean       log2FoldChange             lfcSE
##                        <numeric>            <numeric>         <numeric>
## hsa-mir-206     307.713556384502    -1.33068564558869  0.31505843163435
## hsa-mir-223-5p   459.11883569172   -0.615199198839776 0.169101214163155
## hsa-mir-25-3p   4820.42085197208    0.430107971473196 0.135801218563743
## hsa-mir-10a-5p  346.157958154898   -0.461120240691358 0.156566195466845
## hsa-mir-200a-3p 80.6044955883249    -1.09287077881835 0.370468286286468
## ...                          ...                  ...               ...
## hsa-mir-660-5p  155.677865957598   0.0147634968852977 0.257280472213827
## hsa-mir-92b-3p  64.5712400305805  -0.0213800813149473 0.314517885127516
## hsa-mir-339-5p  107.204731203206  0.00414465840788723 0.266443362717535
## hsa-mir-141-3p  185.044728072799  -0.0010733625846962 0.338906004582831
## hsa-mir-664a-5p 95.2512944422679 -0.00139353456409118 0.282058993696403
##                                 stat               pvalue                padj
##                            <numeric>            <numeric>           <numeric>
## hsa-mir-206        -4.22361540583385 2.40414173602933e-05 0.00548144315814688
## hsa-mir-223-5p     -3.63805311442773 0.000274706780777468  0.0313165730086313
## hsa-mir-25-3p       3.16718786489615  0.00153920848027874   0.116979844501184
## hsa-mir-10a-5p     -2.94520946438279  0.00322736091714395   0.117641431481693
## hsa-mir-200a-3p    -2.94997121014908  0.00317803541903545   0.117641431481693
## ...                              ...                  ...                 ...
## hsa-mir-660-5p    0.0573828894134945    0.954240192811382   0.989272592642855
## hsa-mir-92b-3p   -0.0679773148871299    0.945803692602105   0.989272592642855
## hsa-mir-339-5p    0.0155554950426035     0.98758901119346   0.996328736956234
## hsa-mir-141-3p  -0.00316713947283828    0.997472992537353   0.997472992537353
## hsa-mir-664a-5p -0.00494057837273263    0.996058004831867   0.997472992537353
```

```
mcols(res)$description
```

```
## [1] "mean of normalized counts for all samples"
## [2] "log2 fold change (MLE): condition m vs f" 
## [3] "standard error: condition m vs f"         
## [4] "Wald statistic: condition m vs f"         
## [5] "Wald test p-value: condition m vs f"      
## [6] "BH adjusted p-values"
```

```
# Create a data frame of the ordered results (top 20)
padj.subset <- head(resOrdered, 20) %>%
  as_tibble(rownames = "miR")

# Create a gene table of the subset
#gt(subset)

# Create a interactive gene table of the subset
datatable(padj.subset, 
          extensions = c('KeyTable', "FixedHeader"), 
          caption = 'Table 1: Differentially expressed c-miRs between sexes in path_MMR carriers',
          options = list(keys = TRUE, searchHighlight = TRUE, pageLength = 20, lengthMenu = c("10", "25", "50", "100"))) %>%
  formatRound(columns=c(2:7), digits=3)
```

```
# Create a data frame of all results for plotting
res.df <- as_tibble(res, rownames = "miR")

# Create a volcano plot of results
v1 <- ggplot(data=res.df,
  aes(y=-log10(res$padj), x=res$log2FoldChange, text = paste(rownames(Counts)))) +
  xlab("Log2FC") +
ylab("-log10(Padj)") +
  geom_point(size=2) +
  geom_hline(yintercept = -log10(0.05), linetype="longdash", colour="grey", size=1) +
  geom_vline(xintercept = 1, linetype="longdash", colour="#BE684D", size=1) +
  geom_vline(xintercept = -1, linetype="longdash", colour="#2C467A", size=1) +
  labs(title="Volcano plot",
       subtitle = "Men vs women in LS group",
       caption=paste0("produced on ", Sys.time())) +
  theme_bw()

# Create an interactive volcano plot of results
ggplotly(v1)
```

```
# We saw that there is a sex difference within LS group, so to confirm that this is an unique finding, we need to perform the same analysis to SRME and CTRL group also

# Run the script again from the beginning but subset the groups from phenodata

print("Step 2 completed! There is a sex difference only in LS.")
```

```
## [1] "Step 2 completed! There is a sex difference only in LS."
```

---

#### 5.3 Step 3 - *Path\_MMR* variants in LS

This script is used to perform DE-analysis between MLH1 and other variants (MSH2, MSH6 and PMS2) among *path\_MMR* carriers.

```
# Setup design matrix for DE-analysis
condition <- as.character(Targets$step_3) # condition of interest (path_MMR variant)
batch <- as.character (Targets$NGS) # Batch effect 
sex <- as.character(Targets$Sex) # Sex as covariate
group_levels <- levels(as.factor(condition))
design <- data.frame(condition=as.factor(condition), batch=batch, sex=sex)
rownames(design) <- colnames(Counts)
dds <- DESeqDataSetFromMatrix(countData=Counts, colData=design, design = ~ batch + sex + condition)

# DESeq2 DE-analysis of the condition of interest, batch effect and sex as covariates
dds <- DESeq(dds)

# Display results
res <- results(dds, alpha=0.05)
resOrdered <- res[order(res$padj),]
summary(res)
```

```
## 
## out of 228 with nonzero total read count
## adjusted p-value < 0.05
## LFC > 0 (up)       : 0, 0%
## LFC < 0 (down)     : 0, 0%
## outliers [1]       : 0, 0%
## low counts [2]     : 0, 0%
## (mean count < 8)
## [1] see 'cooksCutoff' argument of ?results
## [2] see 'independentFiltering' argument of ?results
```

```
resOrdered
```

```
## log2 fold change (MLE): condition YES vs NO 
## Wald test p-value: condition YES vs NO 
## DataFrame with 228 rows and 6 columns
##                         baseMean        log2FoldChange             lfcSE
##                        <numeric>             <numeric>         <numeric>
## hsa-mir-206     307.713556384502    -0.918700279522276 0.317688589739701
## hsa-mir-382-5p  480.873186468193     -0.89617112870515 0.300633964174415
## hsa-mir-3065-5p  28.929340476312     0.832735664944019 0.305878893197249
## hsa-mir-155-5p  719.638264871698     0.487418932393798 0.188214918704313
## hsa-mir-185-5p  1172.42293127207    -0.327633576073769 0.132678647051713
## ...                          ...                   ...               ...
## hsa-mir-3613-5p 148.694191498994    0.0126353013852315 0.288019149661537
## hsa-mir-423-5p   7564.1850783196   0.00586543133990573 0.197717936854708
## hsa-mir-664a-5p 95.2512944422679    0.0163241981203909 0.294919727552127
## hsa-mir-20a-5p  872.103417280602 -0.000792319956047674 0.191890092408413
## hsa-mir-361-5p  333.398661677954 -0.000487762461682228 0.131883549541673
##                                 stat              pvalue              padj
##                            <numeric>           <numeric>         <numeric>
## hsa-mir-206        -2.89182649044781 0.00383009405770217 0.436630722578048
## hsa-mir-382-5p     -2.98093773657999 0.00287367213209918 0.436630722578048
## hsa-mir-3065-5p     2.72243585112955 0.00648026131387386 0.492499859854413
## hsa-mir-155-5p      2.58969339810696 0.00960614468691482 0.547550247154145
## hsa-mir-185-5p     -2.46937682403461  0.0135348606646302 0.617189646307139
## ...                              ...                 ...               ...
## hsa-mir-3613-5p    0.043869657278274    0.96500830199003 0.996259041239742
## hsa-mir-423-5p    0.0296656511453279   0.976333706272898 0.996259041239742
## hsa-mir-664a-5p   0.0553513264639296   0.955858572203358 0.996259041239742
## hsa-mir-20a-5p  -0.00412903004059909   0.996705520040721 0.997049084198545
## hsa-mir-361-5p  -0.00369843292341856   0.997049084198545 0.997049084198545
```

```
mcols(res)$description
```

```
## [1] "mean of normalized counts for all samples"  
## [2] "log2 fold change (MLE): condition YES vs NO"
## [3] "standard error: condition YES vs NO"        
## [4] "Wald statistic: condition YES vs NO"        
## [5] "Wald test p-value: condition YES vs NO"     
## [6] "BH adjusted p-values"
```

```
# Create a data frame of the ordered results (top 20)
subset <- head(resOrdered, 20) %>%
  as_tibble(rownames = "miR")

# Create a gene table of the subset
gt(subset)
```

| miR | baseMean | log2FoldChange | lfcSE | stat | pvalue | padj |
| --- | --- | --- | --- | --- | --- | --- |
| hsa-mir-206 | 307.71356 | -0.91870028 | 0.3176886 | -2.8918265 | 0.003830094 | 0.4366307 |
| hsa-mir-382-5p | 480.87319 | -0.89617113 | 0.3006340 | -2.9809377 | 0.002873672 | 0.4366307 |
| hsa-mir-3065-5p | 28.92934 | 0.83273566 | 0.3058789 | 2.7224359 | 0.006480261 | 0.4924999 |
| hsa-mir-155-5p | 719.63826 | 0.48741893 | 0.1882149 | 2.5896934 | 0.009606145 | 0.5475502 |
| hsa-mir-185-5p | 1172.42293 | -0.32763358 | 0.1326786 | -2.4693768 | 0.013534861 | 0.6171896 |
| hsa-mir-22-3p | 785.73921 | -0.36713202 | 0.1549583 | -2.3692306 | 0.017825136 | 0.6773552 |
| hsa-let-7a-5p | 55093.52537 | 0.07330551 | 0.1767418 | 0.4147604 | 0.678317285 | 0.9730676 |
| hsa-let-7c-5p | 1064.66630 | -0.19312682 | 0.1647984 | -1.1718973 | 0.241238279 | 0.9730676 |
| hsa-let-7d-3p | 205.22656 | -0.19022601 | 0.2483986 | -0.7658096 | 0.443789578 | 0.9730676 |
| hsa-let-7f-5p | 27096.20520 | 0.12854418 | 0.1304796 | 0.9851669 | 0.324542076 | 0.9730676 |
| hsa-let-7g-5p | 3442.14965 | -0.09148421 | 0.1776490 | -0.5149718 | 0.606572717 | 0.9730676 |
| hsa-let-7i-5p | 24698.63988 | 0.09293477 | 0.1315752 | 0.7063245 | 0.479986354 | 0.9730676 |
| hsa-mir-101-3p | 1120.60950 | 0.21284126 | 0.2255534 | 0.9436401 | 0.345353602 | 0.9730676 |
| hsa-mir-103a-3p | 2938.41602 | 0.05112773 | 0.1278369 | 0.3999450 | 0.689197034 | 0.9730676 |
| hsa-mir-103b | 1023.43521 | 0.04833881 | 0.1296346 | 0.3728850 | 0.709234031 | 0.9730676 |
| hsa-mir-106b-3p | 206.50278 | -0.28818442 | 0.2205021 | -1.3069465 | 0.191230871 | 0.9730676 |
| hsa-mir-106b-5p | 65.25566 | -0.36266101 | 0.3971767 | -0.9130975 | 0.361191277 | 0.9730676 |
| hsa-mir-107 | 166.46817 | -0.25047445 | 0.1729971 | -1.4478538 | 0.147657941 | 0.9730676 |
| hsa-mir-10b-5p | 394.00082 | -0.03916555 | 0.1469321 | -0.2665555 | 0.789811447 | 0.9730676 |
| hsa-mir-11400 | 46.33394 | -0.14462079 | 0.4377049 | -0.3304070 | 0.741092419 | 0.9730676 |

```
print("Step 3 completed! There was no difference in c-miR expression among path_MMR variants.")
```

```
## [1] "Step 3 completed! There was no difference in c-miR expression among path_MMR variants."
```

---

#### 5.4 Step 4 - Cancer history in *path\_MMR* carriers

This script is used to perform DE-analysis between *path\_MMR carriers* with or without previous cancer(s).

```
# Setup design matrix for DE-analysis
condition <- as.character (Targets$Previous_ca) # Condition of interest (cancer history, 0 = no, 1 = yes)
batch <- as.character (Targets$NGS) # Batch effect
sex <- as.character(Targets$Sex) # Sex as covariate
group_levels <- levels(as.factor(condition))
design <- data.frame(condition=as.factor(condition), batch=batch, sex=sex)
rownames(design) <- colnames(Counts)
dds <- DESeqDataSetFromMatrix(countData=Counts, colData=design, design = ~ batch + sex + condition)

# DESeq2 DE-analysis of the condition of interest, batch effect and sex as covariates
dds <- DESeq(dds)

# Display results
res <- results(dds, alpha=0.05)
resOrdered <- res[order(res$padj),]
summary(res)
```

```
## 
## out of 228 with nonzero total read count
## adjusted p-value < 0.05
## LFC > 0 (up)       : 0, 0%
## LFC < 0 (down)     : 0, 0%
## outliers [1]       : 0, 0%
## low counts [2]     : 0, 0%
## (mean count < 8)
## [1] see 'cooksCutoff' argument of ?results
## [2] see 'independentFiltering' argument of ?results
```

```
resOrdered
```

```
## log2 fold change (MLE): condition 1 vs 0 
## Wald test p-value: condition 1 vs 0 
## DataFrame with 228 rows and 6 columns
##                        baseMean       log2FoldChange             lfcSE
##                       <numeric>            <numeric>         <numeric>
## hsa-mir-206    307.713556384502    0.923234747428217   0.3042997072089
## hsa-mir-140-3p 153.441332464748     0.69732033975536 0.261657008482432
## hsa-mir-224-5p 380.133634872442    0.777869288843377  0.32485293193053
## hsa-mir-375-3p  133.75566338306   -0.790275906805975  0.32494477377035
## hsa-let-7a-5p  55093.5253734347    0.187963807864038 0.168589326071645
## ...                         ...                  ...               ...
## hsa-mir-671-5p 20.9477534559676   0.0198160870856095 0.451436218964841
## hsa-mir-93-5p  4722.36954674415 -0.00739775157996786 0.135343368357135
## hsa-mir-942-5p 47.6943912997957  -0.0763723738706613 0.416039815635458
## hsa-mir-95-3p  31.9363719789938   0.0551554343302459 0.393108998040146
## hsa-mir-30e-3p 165.140350431392  0.00183674883319789 0.263439159317336
##                               stat              pvalue              padj
##                          <numeric>           <numeric>         <numeric>
## hsa-mir-206       3.03396528342507 0.00241362203166191 0.550305823218916
## hsa-mir-140-3p    2.66501686234091  0.0076984462488131 0.877622872364694
## hsa-mir-224-5p    2.39452753041406  0.0166417940627821 0.948582261578581
## hsa-mir-375-3p   -2.43203144225514  0.0150144038690078 0.948582261578581
## hsa-let-7a-5p     1.11492116519975   0.264884205972972 0.969981647919622
## ...                            ...                 ...               ...
## hsa-mir-671-5p  0.0438956518177662    0.96498758130869 0.992728816443445
## hsa-mir-93-5p  -0.0546591360165293   0.956410025400524 0.992728816443445
## hsa-mir-942-5p   -0.18356986759551   0.854350903544122 0.992728816443445
## hsa-mir-95-3p    0.140305703011696   0.888418458546679 0.992728816443445
## hsa-mir-30e-3p 0.00697219364789029   0.994437039404086 0.994437039404086
```

```
mcols(res)$description
```

```
## [1] "mean of normalized counts for all samples"
## [2] "log2 fold change (MLE): condition 1 vs 0" 
## [3] "standard error: condition 1 vs 0"         
## [4] "Wald statistic: condition 1 vs 0"         
## [5] "Wald test p-value: condition 1 vs 0"      
## [6] "BH adjusted p-values"
```

```
# Create a data frame of the ordered results (top 20)
subset <- head(resOrdered, 20) %>%
  as_tibble(rownames = "miR")

# Create a gene table of the subset
gt(subset)
```

| miR | baseMean | log2FoldChange | lfcSE | stat | pvalue | padj |
| --- | --- | --- | --- | --- | --- | --- |
| hsa-mir-206 | 307.71356 | 0.92323475 | 0.3042997 | 3.0339653 | 0.002413622 | 0.5503058 |
| hsa-mir-140-3p | 153.44133 | 0.69732034 | 0.2616570 | 2.6650169 | 0.007698446 | 0.8776229 |
| hsa-mir-224-5p | 380.13363 | 0.77786929 | 0.3248529 | 2.3945275 | 0.016641794 | 0.9485823 |
| hsa-mir-375-3p | 133.75566 | -0.79027591 | 0.3249448 | -2.4320314 | 0.015014404 | 0.9485823 |
| hsa-let-7a-5p | 55093.52537 | 0.18796381 | 0.1685893 | 1.1149212 | 0.264884206 | 0.9699816 |
| hsa-let-7b-5p | 78617.74407 | 0.19400024 | 0.1523492 | 1.2733918 | 0.202879028 | 0.9699816 |
| hsa-let-7c-5p | 1064.66630 | 0.25875637 | 0.1569028 | 1.6491511 | 0.099116681 | 0.9699816 |
| hsa-let-7d-3p | 205.22656 | 0.12760995 | 0.2388988 | 0.5341590 | 0.593231498 | 0.9699816 |
| hsa-let-7d-5p | 1239.02592 | 0.05446379 | 0.1453732 | 0.3746481 | 0.707922232 | 0.9699816 |
| hsa-let-7e-5p | 965.58129 | 0.44854080 | 0.2233915 | 2.0078685 | 0.044657268 | 0.9699816 |
| hsa-let-7f-5p | 27096.20520 | 0.05777348 | 0.1257485 | 0.4594366 | 0.645920641 | 0.9699816 |
| hsa-let-7g-5p | 3442.14965 | -0.06325748 | 0.1705595 | -0.3708822 | 0.710725274 | 0.9699816 |
| hsa-let-7i-5p | 24698.63988 | 0.04418673 | 0.1265056 | 0.3492866 | 0.726874140 | 0.9699816 |
| hsa-mir-100-5p | 46.51049 | -0.27834185 | 0.4416407 | -0.6302450 | 0.528534280 | 0.9699816 |
| hsa-mir-101-3p | 1120.60950 | -0.07714536 | 0.2172718 | -0.3550639 | 0.722541748 | 0.9699816 |
| hsa-mir-103a-3p | 2938.41602 | -0.05941792 | 0.1225922 | -0.4846795 | 0.627903703 | 0.9699816 |
| hsa-mir-103b | 1023.43521 | -0.07940509 | 0.1241771 | -0.6394506 | 0.522529845 | 0.9699816 |
| hsa-mir-106b-3p | 206.50278 | -0.06491278 | 0.2136906 | -0.3037700 | 0.761303126 | 0.9699816 |
| hsa-mir-106b-5p | 65.25566 | -0.15376889 | 0.3825078 | -0.4020020 | 0.687682575 | 0.9699816 |
| hsa-mir-10a-5p | 346.15796 | 0.06618768 | 0.1569210 | 0.4217899 | 0.673178350 | 0.9699816 |

```
print("Step 4 completed! There was no difference in c-miR expression between path_MMR carriers with or without cancer history.")
```

```
## [1] "Step 4 completed! There was no difference in c-miR expression between path_MMR carriers with or without cancer history."
```

---

#### 5.5 Step 5 - Healthy *path\_MMR* carriers vs *path\_MMR* carriers with cancer

This script is used to perform DE-analysis between healthy *path\_MMR* carriers (n=81) and *path\_MMR* carriers with cancer (n=13).

```
# Choose conditions of interest (healthy path_MMR carriers, n = 81 and path_MMR carriers with cancer, n = 13)
select <- which(targets$Type=="LS" & targets$Healthy_now=="YES" | targets$Healthy_now=="NO")

# Create a new filtered counts file
Counts <- counts[,select]

# New phenofile with only the variables of interest
Targets <- targets[select,]

# Setup design matrix for DE-analysis
condition <- as.character (Targets$Healthy_now) # Condition of interest
batch <- as.character (Targets$NGS) # Batch effect
sex <- as.character(Targets$Sex) # Sex as covariate
group_levels <- levels(as.factor(condition))
design <- data.frame(condition=as.factor(condition), batch=batch, sex=sex)
rownames(design) <- colnames(Counts)
dds <- DESeqDataSetFromMatrix(countData=Counts, colData=design, design = ~ batch + sex + condition)

# DESeq2 DE-analysis of the condition of interest, batch effect and sex as covariates
dds <- DESeq(dds)

# Display results
res <- results(dds, alpha=0.05)
resOrdered <- res[order(res$padj),]
summary(res)
```

```
## 
## out of 228 with nonzero total read count
## adjusted p-value < 0.05
## LFC > 0 (up)       : 0, 0%
## LFC < 0 (down)     : 0, 0%
## outliers [1]       : 0, 0%
## low counts [2]     : 0, 0%
## (mean count < 8)
## [1] see 'cooksCutoff' argument of ?results
## [2] see 'independentFiltering' argument of ?results
```

```
resOrdered
```

```
## log2 fold change (MLE): condition YES vs NO 
## Wald test p-value: condition YES vs NO 
## DataFrame with 228 rows and 6 columns
##                         baseMean        log2FoldChange             lfcSE
##                        <numeric>             <numeric>         <numeric>
## hsa-mir-127-3p  48.0155927505647      1.54757991082434 0.478860956403904
## hsa-let-7b-5p   82367.6769607343    -0.250486443035102 0.217244307939718
## hsa-let-7c-5p   1046.64340989249     0.190973049839557  0.21370151185407
## hsa-let-7d-3p   209.630058542978    -0.145838717995448 0.309838889110817
## hsa-let-7d-5p   1343.69887604826    -0.324719189652028 0.221464701451058
## ...                          ...                   ...               ...
## hsa-mir-484     448.505322877159   0.00216435004940099 0.396810802499483
## hsa-mir-503-5p  32.0124906562595     -0.01357393058526 0.505835299507598
## hsa-mir-652-3p  47.3776761888421 -0.000604709085427502  0.48879803619999
## hsa-mir-664a-5p 96.3727540555979 -0.000729931278471826 0.364405578438736
## hsa-mir-99a-5p  101.327988540091   0.00775182190537004 0.420844743177602
##                                 stat              pvalue              padj
##                            <numeric>           <numeric>         <numeric>
## hsa-mir-127-3p       3.2317938853194 0.00123015753681806 0.280475918394519
## hsa-let-7b-5p      -1.15301728920146   0.248903288182279 0.998288877191795
## hsa-let-7c-5p      0.893643887601349    0.37151245670584 0.998288877191795
## hsa-let-7d-3p     -0.470692101995265   0.637860625298168 0.998288877191795
## hsa-let-7d-5p      -1.46623451739459   0.142584407997209 0.998288877191795
## ...                              ...                 ...               ...
## hsa-mir-484      0.00545436272341352   0.995648069772354 0.999012909454964
## hsa-mir-503-5p   -0.0268346843300051   0.978591589077029 0.999012909454964
## hsa-mir-652-3p  -0.00123713485047654   0.999012909454964 0.999012909454964
## hsa-mir-664a-5p -0.00200307383218213   0.998401779383912 0.999012909454964
## hsa-mir-99a-5p    0.0184196714608805   0.985304059547971 0.999012909454964
```

```
mcols(res)$description
```

```
## [1] "mean of normalized counts for all samples"  
## [2] "log2 fold change (MLE): condition YES vs NO"
## [3] "standard error: condition YES vs NO"        
## [4] "Wald statistic: condition YES vs NO"        
## [5] "Wald test p-value: condition YES vs NO"     
## [6] "BH adjusted p-values"
```

```
# Create a data frame of the ordered results (top 20)
subset <- head(resOrdered, 20) %>%
  as_tibble(rownames = "miR")

# Create a gene table of the subset
gt(subset)
```

| miR | baseMean | log2FoldChange | lfcSE | stat | pvalue | padj |
| --- | --- | --- | --- | --- | --- | --- |
| hsa-mir-127-3p | 48.01559 | 1.54757991 | 0.4788610 | 3.2317939 | 0.001230158 | 0.2804759 |
| hsa-let-7b-5p | 82367.67696 | -0.25048644 | 0.2172443 | -1.1530173 | 0.248903288 | 0.9982889 |
| hsa-let-7c-5p | 1046.64341 | 0.19097305 | 0.2137015 | 0.8936439 | 0.371512457 | 0.9982889 |
| hsa-let-7d-3p | 209.63006 | -0.14583872 | 0.3098389 | -0.4706921 | 0.637860625 | 0.9982889 |
| hsa-let-7d-5p | 1343.69888 | -0.32471919 | 0.2214647 | -1.4662345 | 0.142584408 | 0.9982889 |
| hsa-let-7e-5p | 926.64673 | 0.33109997 | 0.2942949 | 1.1250620 | 0.260562767 | 0.9982889 |
| hsa-let-7f-5p | 26808.59858 | 0.17400238 | 0.1598255 | 1.0887025 | 0.276285084 | 0.9982889 |
| hsa-let-7i-5p | 24851.67228 | 0.06746493 | 0.1650282 | 0.4088084 | 0.682680275 | 0.9982889 |
| hsa-mir-100-5p | 48.67038 | -0.45345588 | 0.5825199 | -0.7784384 | 0.436310596 | 0.9982889 |
| hsa-mir-101-3p | 1147.85281 | 0.17065168 | 0.2953940 | 0.5777086 | 0.563460858 | 0.9982889 |
| hsa-mir-103a-3p | 2918.11081 | 0.24630999 | 0.1736895 | 1.4181055 | 0.156159970 | 0.9982889 |
| hsa-mir-103b | 1015.61076 | 0.23920375 | 0.1725654 | 1.3861625 | 0.165697280 | 0.9982889 |
| hsa-mir-106b-3p | 215.96717 | -0.21582880 | 0.2886214 | -0.7477920 | 0.454585615 | 0.9982889 |
| hsa-mir-106b-5p | 66.96291 | 0.53460596 | 0.5182629 | 1.0315342 | 0.302290377 | 0.9982889 |
| hsa-mir-107 | 168.30275 | 0.24345940 | 0.2546063 | 0.9562190 | 0.338961590 | 0.9982889 |
| hsa-mir-10a-5p | 347.95892 | -0.19387889 | 0.2094819 | -0.9255162 | 0.354697482 | 0.9982889 |
| hsa-mir-10b-5p | 400.29913 | -0.17038572 | 0.1877073 | -0.9077203 | 0.364026003 | 0.9982889 |
| hsa-mir-11400 | 43.71408 | 0.54947799 | 0.5573913 | 0.9858029 | 0.324229849 | 0.9982889 |
| hsa-mir-1180-3p | 48.65138 | -0.86372484 | 0.5626079 | -1.5352164 | 0.124730683 | 0.9982889 |
| hsa-mir-122b-3p | 12230.59670 | 0.05827116 | 0.4349263 | 0.1339794 | 0.893418872 | 0.9982889 |

```
print("Step 5 completed! There was no difference in c-miR expression between path_MMR carriers with or without cancer.")
```

```
## [1] "Step 5 completed! There was no difference in c-miR expression between path_MMR carriers with or without cancer."
```

---

#### 5.6 Step 6 - Healthy *path\_MMR* carriers vs non-LS control group

This script is used to perform DE-analysis between healthy *path\_MMR* carriers and CTRL group.

```
# Choose conditions of interest (healthy path_MMR carriers, n = 81 and CTRL group, n = 37)
select <- which(targets$Type=="LS" & targets$Healthy_now=="YES" | targets$Type=="CTRL")

# Create a new filtered counts file
Counts <- counts[,select]

# New phenofile with only the variables of interest
Targets <- targets[select,]

# Setup design matrix for DE-analysis
condition <- as.character (Targets$Type) # Condition of interest
batch <- as.character (Targets$NGS) # Batch effect
sex <- as.character(Targets$Sex) # Sex as covariate
group_levels <- levels(as.factor(condition))
design <- data.frame(condition=as.factor(condition), batch=batch, sex=sex)
rownames(design) <- colnames(Counts)
dds <- DESeqDataSetFromMatrix(countData=Counts, colData=design, design = ~ batch + sex + condition)

# DESeq2 DE-analysis of the condition of interest, batch effect and sex as covariates
dds <- DESeq(dds)

# Display results
res <- results(dds, alpha=0.05)
resOrdered <- res[order(res$padj),]
summary(res)
```

```
## 
## out of 228 with nonzero total read count
## adjusted p-value < 0.05
## LFC > 0 (up)       : 15, 6.6%
## LFC < 0 (down)     : 25, 11%
## outliers [1]       : 0, 0%
## low counts [2]     : 75, 33%
## (mean count < 69)
## [1] see 'cooksCutoff' argument of ?results
## [2] see 'independentFiltering' argument of ?results
```

```
resOrdered
```

```
## log2 fold change (MLE): condition LS vs CTRL 
## Wald test p-value: condition LS vs CTRL 
## DataFrame with 228 rows and 6 columns
##                         baseMean     log2FoldChange             lfcSE
##                        <numeric>          <numeric>         <numeric>
## hsa-mir-155-5p  649.461599394047  0.905013618668636 0.169010672683695
## hsa-let-7c-5p   978.123000672281  0.728728381283735 0.145105235191548
## hsa-let-7e-5p   859.891695461067  0.955495744987595 0.195765328042533
## hsa-mir-122b-3p 10449.9542595749   1.25212835907867 0.293791801308695
## hsa-mir-15a-5p   696.18084962528 -0.677170624079328 0.162771620286646
## ...                          ...                ...               ...
## hsa-mir-769-5p  29.9692330488429 0.0653055737807121 0.340804461425901
## hsa-mir-942-5p  54.8480155832624 -0.591235768488896 0.325881717208209
## hsa-mir-95-3p    29.533154889069  0.691400207993933 0.351805136918705
## hsa-mir-4742-3p 17.8527070960152 0.0262833960823129 0.472512510966432
## hsa-mir-532-3p  23.5287803958615 -0.248081655266415 0.484669483704653
##                               stat               pvalue                 padj
##                          <numeric>            <numeric>            <numeric>
## hsa-mir-155-5p    5.35477200521163 8.56643641079564e-08 1.31066477085173e-05
## hsa-let-7c-5p     5.02206815847662 5.11180354684818e-07 3.91052971333886e-05
## hsa-let-7e-5p      4.8808221279102 1.05644496571581e-06 5.38786932515063e-05
## hsa-mir-122b-3p   4.26195814008787 2.02643407595029e-05 0.000775111034050985
## hsa-mir-15a-5p   -4.16024994336734 3.17899503523146e-05 0.000972772480780828
## ...                            ...                  ...                  ...
## hsa-mir-769-5p   0.191621827682297    0.848038443336644                   NA
## hsa-mir-942-5p   -1.81426492272701   0.0696369491588198                   NA
## hsa-mir-95-3p     1.96529309961071   0.0493803237896099                   NA
## hsa-mir-4742-3p  0.055624762249269    0.955640737618766                   NA
## hsa-mir-532-3p  -0.511857386543425    0.608750823822992                   NA
```

```
mcols(res)$description
```

```
## [1] "mean of normalized counts for all samples"   
## [2] "log2 fold change (MLE): condition LS vs CTRL"
## [3] "standard error: condition LS vs CTRL"        
## [4] "Wald statistic: condition LS vs CTRL"        
## [5] "Wald test p-value: condition LS vs CTRL"     
## [6] "BH adjusted p-values"
```

```
# Create a data frame of the ordered results (top 39)
padj.subset <- head(resOrdered, 40) %>%
 as_tibble(rownames = "miR")

# Create a interactive gene table of the subset
datatable(padj.subset, 
          extensions = c('KeyTable', "FixedHeader"), 
          caption = 'Table 2: Differentially expressed c-miRs in healthy path_MMR carriers vs non-LS control group',
          options = list(keys = TRUE, searchHighlight = TRUE, pageLength = 20, lengthMenu = c("5", "10", "15", "50"))) %>%
  formatRound(columns=c(2:7), digits=3)
```

```
# Create a data frame of all results for plotting
res.df <- as_tibble(res, rownames = "miR")

# Create a volcano plot of results
v2 <- ggplot(data=res.df,
  aes(y=-log10(res$padj), x=res$log2FoldChange, text = paste(rownames(Counts)))) +
  xlab("Log2FC") +
ylab("-log10(Padj)") +
  geom_point(size=2) +
  geom_hline(yintercept = -log10(0.05), linetype="longdash", colour="grey", size=1) +
  geom_vline(xintercept = 1, linetype="longdash", colour="#BE684D", size=1) +
  geom_vline(xintercept = -1, linetype="longdash", colour="#2C467A", size=1) +
  labs(title="Volcano plot",
       subtitle = "LS healthy vs CTRL",
       caption=paste0("produced on ", Sys.time())) +
  theme_bw()

# Create an interactive volcano plot of results
ggplotly(v2)
```

```
print("Step 6 completed! There are 15 upregulated and 25 downregulated c-miRs in healthy path_MMR carriers compared to CTRL group.")
```

```
## [1] "Step 6 completed! There are 15 upregulated and 25 downregulated c-miRs in healthy path_MMR carriers compared to CTRL group."
```

---

#### 5.7 Step 7 - Sporadic rectal cancer patients vs healthy *path\_MMR* carriers

This script is used to perform DE-analysis between SRME group and healthy *path\_MMR* carriers.

```
# Choose conditions of interest (healthy path_MMR carriers, n = 81 and SRME, n = 24)
select <- which(targets$Type=="LS" & targets$Healthy_now=="YES" | targets$Type=="SRME")

# Create a new filtered counts file
Counts <- counts[,select]

# New phenofile with only the variables of interest
Targets <- targets[select,]

# Setup design matrix for DE-analysis
condition <- as.character (Targets$Type) # Condition of interest
batch <- as.character (Targets$NGS) # Batch effect
sex <- as.character(Targets$Sex) # Sex as covariate
group_levels <- levels(as.factor(condition))
design <- data.frame(condition=as.factor(condition), batch=batch, sex=sex)
rownames(design) <- colnames(Counts)
dds <- DESeqDataSetFromMatrix(countData=Counts, colData=design, design = ~ batch + sex + condition)

# DESeq2 DE-analysis of the condition of interest, batch effect and sex as covariates
dds <- DESeq(dds)

# Display results
res <- results(dds, alpha=0.05)
resOrdered <- res[order(res$padj),]
summary(res)
```

```
## 
## out of 228 with nonzero total read count
## adjusted p-value < 0.05
## LFC > 0 (up)       : 0, 0%
## LFC < 0 (down)     : 0, 0%
## outliers [1]       : 0, 0%
## low counts [2]     : 0, 0%
## (mean count < 9)
## [1] see 'cooksCutoff' argument of ?results
## [2] see 'independentFiltering' argument of ?results
```

```
resOrdered
```

```
## log2 fold change (MLE): condition SRME vs LS 
## Wald test p-value: condition SRME vs LS 
## DataFrame with 228 rows and 6 columns
##                         baseMean       log2FoldChange             lfcSE
##                        <numeric>            <numeric>         <numeric>
## hsa-mir-10a-5p  415.729129524597    0.700061546934253 0.197350126933255
## hsa-mir-1180-3p 55.3191753024436      1.1469344634644 0.407027948054128
## hsa-mir-126-3p  27858.2897731659   -0.395198316713981 0.136574204015264
## hsa-mir-148b-3p 1429.34443333521   -0.335860304942317 0.111582302782957
## hsa-mir-196a-5p 32.2507910563298     1.41409583370707 0.497316069110564
## ...                          ...                  ...               ...
## hsa-mir-152-3p  101.315350736564  -0.0117083365991052 0.300777798273116
## hsa-mir-15a-5p  565.801351946751 -0.00508602975432469  0.19247823226543
## hsa-mir-19a-3p  200.176285402756  0.00281087665664761 0.231068789963995
## hsa-mir-20b-5p  118.924674916842  0.00375727754095451 0.266008834305326
## hsa-mir-485-3p  46.0863537963314   0.0134266672579918 0.608534438893608
##                                stat               pvalue               padj
##                           <numeric>            <numeric>          <numeric>
## hsa-mir-10a-5p     3.54730730510762 0.000389190337738628 0.0887353970044071
## hsa-mir-1180-3p     2.8178272989546  0.00483498082966978  0.157482232737816
## hsa-mir-126-3p    -2.89365271841388  0.00380789063706925  0.157482232737816
## hsa-mir-148b-3p   -3.00997825430804  0.00261266394498759  0.157482232737816
## hsa-mir-196a-5p    2.84345494050924   0.0044627319143887  0.157482232737816
## ...                             ...                  ...                ...
## hsa-mir-152-3p  -0.0389268645037214    0.968948698020339   0.99029423326846
## hsa-mir-15a-5p  -0.0264239217830668    0.978919213985837   0.99029423326846
## hsa-mir-19a-3p   0.0121646746714932     0.99029423326846   0.99029423326846
## hsa-mir-20b-5p   0.0141246344346665    0.988730546978105   0.99029423326846
## hsa-mir-485-3p   0.0220639398526124    0.982396951297894   0.99029423326846
```

```
mcols(res)$description
```

```
## [1] "mean of normalized counts for all samples"   
## [2] "log2 fold change (MLE): condition SRME vs LS"
## [3] "standard error: condition SRME vs LS"        
## [4] "Wald statistic: condition SRME vs LS"        
## [5] "Wald test p-value: condition SRME vs LS"     
## [6] "BH adjusted p-values"
```

```
# Create a data frame of the ordered results (top 20)
subset<- head(resOrdered, 20) %>%
  as_tibble(rownames = "miR")

# Create a gene table of the subset
gt(subset)
```

| miR | baseMean | log2FoldChange | lfcSE | stat | pvalue | padj |
| --- | --- | --- | --- | --- | --- | --- |
| hsa-mir-10a-5p | 415.72913 | 0.7000615 | 0.1973501 | 3.547307 | 0.0003891903 | 0.0887354 |
| hsa-mir-1180-3p | 55.31918 | 1.1469345 | 0.4070279 | 2.817827 | 0.0048349808 | 0.1574822 |
| hsa-mir-126-3p | 27858.28977 | -0.3951983 | 0.1365742 | -2.893653 | 0.0038078906 | 0.1574822 |
| hsa-mir-148b-3p | 1429.34443 | -0.3358603 | 0.1115823 | -3.009978 | 0.0026126639 | 0.1574822 |
| hsa-mir-196a-5p | 32.25079 | 1.4140958 | 0.4973161 | 2.843455 | 0.0044627319 | 0.1574822 |
| hsa-mir-320a-3p | 1260.89967 | 0.5573208 | 0.1969468 | 2.829804 | 0.0046576534 | 0.1574822 |
| hsa-mir-320b | 138.33790 | 0.8445221 | 0.2851437 | 2.961742 | 0.0030590444 | 0.1574822 |
| hsa-mir-486-5p | 28352.51084 | 0.5415406 | 0.2062100 | 2.626161 | 0.0086354078 | 0.2461091 |
| hsa-mir-320c | 23.21182 | 1.0968594 | 0.4285908 | 2.559223 | 0.0104906547 | 0.2657633 |
| hsa-mir-185-5p | 1266.34806 | 0.3435518 | 0.1399213 | 2.455321 | 0.0140758737 | 0.2917545 |
| hsa-mir-223-3p | 36332.64595 | 0.4133965 | 0.1661891 | 2.487507 | 0.0128641845 | 0.2917545 |
| hsa-mir-483-5p | 260.32253 | 0.7739761 | 0.3242560 | 2.386929 | 0.0169897539 | 0.3228053 |
| hsa-mir-2110 | 19.66771 | 1.1984973 | 0.5204457 | 2.302829 | 0.0212884818 | 0.3466981 |
| hsa-mir-222-3p | 117.27440 | 0.7502785 | 0.3226651 | 2.325255 | 0.0200583342 | 0.3466981 |
| hsa-mir-486-3p | 297.90130 | 0.4754586 | 0.2164429 | 2.196693 | 0.0280423622 | 0.4262439 |
| hsa-let-7d-3p | 227.49748 | 0.4615138 | 0.2285625 | 2.019201 | 0.0434662854 | 0.4677974 |
| hsa-mir-11400 | 42.63983 | -0.8238021 | 0.4118758 | -2.000123 | 0.0454870007 | 0.4677974 |
| hsa-mir-134-5p | 120.39936 | -0.6698154 | 0.3319645 | -2.017732 | 0.0436192208 | 0.4677974 |
| hsa-mir-193a-5p | 94.50090 | 0.5318262 | 0.2682622 | 1.982487 | 0.0474247630 | 0.4677974 |
| hsa-mir-196b-5p | 316.86277 | 0.4468163 | 0.2272149 | 1.966492 | 0.0492418312 | 0.4677974 |

```
print("Step 7 completed! There was no difference in c-miR expression between SRME group and healthy path_MMR carriers.")
```

```
## [1] "Step 7 completed! There was no difference in c-miR expression between SRME group and healthy path_MMR carriers."
```

---

#### 5.8 Step 8 - Sporadic rectal cancer patients vs *path\_MMR* carriers with cancer

This script is used to perform DE-analysis between SRME group and *path\_MMR* carriers with cancer.

```
# Choose conditions of interest (path_MMR with cancer, n = 13 and SRME, n = 24)
select <- which(targets$Type=="LS" & targets$Healthy_now=="NO" | targets$Type=="SRME")

# Create a new filtered counts file
Counts <- counts[,select]

# New phenofile with only the variables of interest
Targets <- targets[select,]

# Setup design matrix for DE-analysis
condition <- as.character (Targets$Type) # Condition of interest
batch <- as.character (Targets$NGS) # Batch effect
sex <- as.character(Targets$Sex) # Sex as covariate
group_levels <- levels(as.factor(condition))
design <- data.frame(condition=as.factor(condition), batch=batch, sex=sex)
rownames(design) <- colnames(Counts)
dds <- DESeqDataSetFromMatrix(countData=Counts, colData=design, design = ~ batch + sex + condition)

# DESeq2 DE-analysis of the condition of interest, batch effect and sex as covariates
dds <- DESeq(dds)

# Display results
res <- results(dds, alpha=0.05)
resOrdered <- res[order(res$padj),]
summary(res)
```

```
## 
## out of 228 with nonzero total read count
## adjusted p-value < 0.05
## LFC > 0 (up)       : 0, 0%
## LFC < 0 (down)     : 0, 0%
## outliers [1]       : 0, 0%
## low counts [2]     : 0, 0%
## (mean count < 9)
## [1] see 'cooksCutoff' argument of ?results
## [2] see 'independentFiltering' argument of ?results
```

```
resOrdered
```

```
## log2 fold change (MLE): condition SRME vs LS 
## Wald test p-value: condition SRME vs LS 
## DataFrame with 228 rows and 6 columns
##                         baseMean      log2FoldChange             lfcSE
##                        <numeric>           <numeric>         <numeric>
## hsa-let-7a-5p   60333.0739919861  0.0092484018664266 0.292951248812435
## hsa-let-7b-5p    101740.53065124  0.0649964919100044 0.278744679867905
## hsa-let-7c-5p   1191.71906684258   0.222884217723653 0.313235279614099
## hsa-let-7d-3p   277.106598697734   0.295007448481522 0.257524716257159
## hsa-let-7d-5p   1620.79876824715  -0.229396853580775 0.258804980000646
## ...                          ...                 ...               ...
## hsa-mir-99a-5p  111.185435654031   0.393622024916593 0.546722550230911
## hsa-mir-99b-5p    136.2175515144  0.0391449465983428 0.364010303946211
## hsa-mir-4742-3p 24.7866960150387    1.67461261850616  0.91933989739968
## hsa-mir-532-3p  35.3127536910979   -0.20117446613601 0.758781908425807
## hsa-mir-30e-5p  3943.14710049544 0.00187342365499215 0.227162529976706
##                                stat             pvalue              padj
##                           <numeric>          <numeric>         <numeric>
## hsa-let-7a-5p    0.0315697642659581  0.974815155986009 0.984058113657596
## hsa-let-7b-5p     0.233175721742226  0.815624957520626 0.984058113657596
## hsa-let-7c-5p     0.711555282017541  0.476740206505712 0.984058113657596
## hsa-let-7d-3p      1.14555003795027  0.251981380053937 0.984058113657596
## hsa-let-7d-5p     -0.88636954969028  0.375418422243555 0.984058113657596
## ...                             ...                ...               ...
## hsa-mir-99a-5p    0.719966690143557  0.471545504767019 0.984058113657596
## hsa-mir-99b-5p    0.107538017946127  0.914362166220587 0.984058113657596
## hsa-mir-4742-3p    1.82153806578257 0.0685251057836261 0.984058113657596
## hsa-mir-532-3p   -0.265128179654906   0.79091071965623 0.984058113657596
## hsa-mir-30e-5p  0.00824706281966599  0.993419870494506 0.993419870494506
```

```
mcols(res)$description
```

```
## [1] "mean of normalized counts for all samples"   
## [2] "log2 fold change (MLE): condition SRME vs LS"
## [3] "standard error: condition SRME vs LS"        
## [4] "Wald statistic: condition SRME vs LS"        
## [5] "Wald test p-value: condition SRME vs LS"     
## [6] "BH adjusted p-values"
```

```
# Create a data frame of the ordered results (top 20)
subset<- head(resOrdered, 20) %>%
  as_tibble(rownames = "miR")

# Create a gene table of the subset
gt(subset)
```

| miR | baseMean | log2FoldChange | lfcSE | stat | pvalue | padj |
| --- | --- | --- | --- | --- | --- | --- |
| hsa-let-7a-5p | 60333.07399 | 0.009248402 | 0.2929512 | 0.03156976 | 0.97481516 | 0.9840581 |
| hsa-let-7b-5p | 101740.53065 | 0.064996492 | 0.2787447 | 0.23317572 | 0.81562496 | 0.9840581 |
| hsa-let-7c-5p | 1191.71907 | 0.222884218 | 0.3132353 | 0.71155528 | 0.47674021 | 0.9840581 |
| hsa-let-7d-3p | 277.10660 | 0.295007448 | 0.2575247 | 1.14555004 | 0.25198138 | 0.9840581 |
| hsa-let-7d-5p | 1620.79877 | -0.229396854 | 0.2588050 | -0.88636955 | 0.37541842 | 0.9840581 |
| hsa-let-7e-5p | 904.85522 | 0.260591230 | 0.3587440 | 0.72639881 | 0.46759432 | 0.9840581 |
| hsa-let-7f-5p | 28049.73885 | 0.236805612 | 0.2387517 | 0.99184886 | 0.32127126 | 0.9840581 |
| hsa-let-7g-5p | 4425.48657 | 0.194992567 | 0.2452425 | 0.79510113 | 0.42655468 | 0.9840581 |
| hsa-let-7i-5p | 25755.91770 | -0.109233089 | 0.2233584 | -0.48904839 | 0.62480744 | 0.9840581 |
| hsa-mir-100-5p | 56.87738 | -0.585330355 | 0.5831189 | -1.00379246 | 0.31547866 | 0.9840581 |
| hsa-mir-101-3p | 1060.15111 | -0.255158402 | 0.3007912 | -0.84829073 | 0.39627608 | 0.9840581 |
| hsa-mir-103a-3p | 2860.81663 | 0.135676208 | 0.2676541 | 0.50690881 | 0.61221880 | 0.9840581 |
| hsa-mir-103b | 985.95365 | 0.107224271 | 0.2608018 | 0.41113317 | 0.68097489 | 0.9840581 |
| hsa-mir-106b-3p | 254.61759 | -0.592788554 | 0.3516875 | -1.68555492 | 0.09188156 | 0.9840581 |
| hsa-mir-106b-5p | 81.11799 | 0.459848128 | 0.5790011 | 0.79420938 | 0.42707355 | 0.9840581 |
| hsa-mir-107 | 172.92029 | 0.262385177 | 0.3688201 | 0.71141780 | 0.47682537 | 0.9840581 |
| hsa-mir-10a-5p | 563.55545 | 0.283853049 | 0.4081588 | 0.69544761 | 0.48677483 | 0.9840581 |
| hsa-mir-10b-5p | 465.30640 | 0.142811036 | 0.2315570 | 0.61674234 | 0.53740469 | 0.9840581 |
| hsa-mir-11400 | 29.39506 | -0.085038750 | 0.5911966 | -0.14384174 | 0.88562544 | 0.9840581 |
| hsa-mir-1180-3p | 88.54225 | 0.075759366 | 0.4876652 | 0.15535118 | 0.87654447 | 0.9840581 |

```
print("Step 8 completed! There was no difference in c-miR expression between SRME group and path_MMR carriers with cancer.")
```

```
## [1] "Step 8 completed! There was no difference in c-miR expression between SRME group and path_MMR carriers with cancer."
```

---

#### 5.9 Step 9 - Sporadic rectal cancer patients vs non-LS control group

This script is used to perform DE-analysis between SRME group and CTRL group.

```
# Choose conditions of interest (SRME, n = 24, CTRL, n = 37)
select <- which(targets$Type=="SRME" | targets$Type=="CTRL")

# Create a new filtered counts file
Counts <- counts[,select]

# New phenofile with only the variables of interest
Targets <- targets[select,]

# Setup design matrix for DE-analysis
condition <- as.character (Targets$Type) # Condition of interest
batch <- as.character (Targets$NGS) # Batch effect
sex <- as.character(Targets$Sex) # Sex as covariate
group_levels <- levels(as.factor(condition))
design <- data.frame(condition=as.factor(condition), batch=batch, sex=sex)
rownames(design) <- colnames(Counts)
dds <- DESeqDataSetFromMatrix(countData=Counts, colData=design, design = ~ batch + sex + condition)

# DESeq2 DE-analysis of the condition of interest, batch effect and sex as covariates
dds <- DESeq(dds)

# Display results
res <- results(dds, alpha=0.05)
resOrdered <- res[order(res$padj),]
summary(res)
```

```
## 
## out of 228 with nonzero total read count
## adjusted p-value < 0.05
## LFC > 0 (up)       : 4, 1.8%
## LFC < 0 (down)     : 0, 0%
## outliers [1]       : 0, 0%
## low counts [2]     : 0, 0%
## (mean count < 9)
## [1] see 'cooksCutoff' argument of ?results
## [2] see 'independentFiltering' argument of ?results
```

```
resOrdered
```

```
## log2 fold change (MLE): condition SRME vs CTRL 
## Wald test p-value: condition SRME vs CTRL 
## DataFrame with 228 rows and 6 columns
##                         baseMean       log2FoldChange             lfcSE
##                        <numeric>            <numeric>         <numeric>
## hsa-mir-200a-3p 111.715251574488     1.75586404983053  0.37754305822285
## hsa-mir-10a-5p   466.81555503228    0.980500751171361 0.231783050278878
## hsa-mir-196a-5p 37.3750252312532     1.81268787183865 0.510932292180228
## hsa-mir-200c-3p 124.838948446616     1.13256759077687 0.324663822253165
## hsa-let-7e-5p   750.356684894863    0.721450941450189 0.236915042105674
## ...                          ...                  ...               ...
## hsa-let-7d-5p   1568.65433721735 -0.00574460659022131 0.148270261107422
## hsa-mir-3065-5p 33.7577867286205  0.00948004175628501  0.33179298733574
## hsa-mir-382-5p   466.84694089799  0.00765254387100355 0.299528556713819
## hsa-mir-106b-5p 93.2843324209975 -0.00414373615461178 0.352729339070495
## hsa-mir-4732-3p  36.556860995693  0.00312193397737304 0.431178830243722
##                                stat               pvalue                 padj
##                           <numeric>            <numeric>            <numeric>
## hsa-mir-200a-3p    4.65076502292383 3.30705960597195e-06 0.000754009590161605
## hsa-mir-10a-5p     4.23025216896419 2.33429489844175e-05  0.00266109618422359
## hsa-mir-196a-5p    3.54780447347265 0.000388456479126744    0.027694180338899
## hsa-mir-200c-3p    3.48843176587048 0.000485862812963141    0.027694180338899
## hsa-let-7e-5p      3.04518841453888  0.00232534584517635   0.0920451642655798
## ...                             ...                  ...                  ...
## hsa-let-7d-5p   -0.0387441591274959    0.969094365929624    0.986399622464081
## hsa-mir-3065-5p  0.0285721582978853    0.977205817475231    0.988286541248025
## hsa-mir-382-5p    0.025548628668201    0.979617361061639    0.988286541248025
## hsa-mir-106b-5p -0.0117476367730886    0.990626957584192    0.994222997555712
## hsa-mir-4732-3p 0.00724046209691783    0.994222997555712    0.994222997555712
```

```
mcols(res)$description
```

```
## [1] "mean of normalized counts for all samples"     
## [2] "log2 fold change (MLE): condition SRME vs CTRL"
## [3] "standard error: condition SRME vs CTRL"        
## [4] "Wald statistic: condition SRME vs CTRL"        
## [5] "Wald test p-value: condition SRME vs CTRL"     
## [6] "BH adjusted p-values"
```

```
# Create a data frame of the ordered results (top 3)
padj.subset<- head(resOrdered, 4) %>%
  as_tibble(rownames = "miR")

# Create a interactive gene table of the subset
datatable(padj.subset, 
          extensions = c('KeyTable', "FixedHeader"), 
          caption = 'Table 3: Differentially expressed c-miRs in sporadic rectal cancer patients vs non-LS control group',
          options = list(keys = TRUE, searchHighlight = TRUE, pageLength = 20, lengthMenu = c("5", "10", "15", "50"))) %>%
  formatRound(columns=c(2:7), digits=3)
```

```
# Create a data frame of all results for plotting
res.df <- as_tibble(res, rownames = "miR")

# Create a volcano plot of results
v3 <- ggplot(data=res.df,
             aes(y=-log10(res$padj), x=res$log2FoldChange)) +
  xlab("Log2FC") +
  ylab("-log10(Padj)") +
  geom_point(size=2) +
  geom_hline(yintercept = -log10(0.05), linetype="longdash", colour="grey", size=1) +
  geom_vline(xintercept = 1, linetype="longdash", colour="#BE684D", size=1) +
  geom_vline(xintercept = -1, linetype="longdash", colour="#2C467A", size=1) +
  labs(title="Volcano plot",
       subtitle = "LS vs SRME",
       caption=paste0("produced on ", Sys.time())) +
  theme_bw()

ggplotly(v3)
```

```
print("Step 9 completed! There were 4 upregulated c-miRs in SRME group compared to CTRL.")
```

```
## [1] "Step 9 completed! There were 4 upregulated c-miRs in SRME group compared to CTRL."
```

---

#### 5.10 Step 10 - *Path\_MMR* carriers with cancer vs non-LS control group

This script is used to perform DE-analysis between *path\_MMR* carriers with cancer and CTRL group.

```
# Choose conditions of interest (Path_MMR carriers with cancer, n = 13 and CTRL, n = 37)
select <- which(targets$Type=="LS" & targets$Healthy_now=="NO" | targets$Type=="CTRL")

# Create a new filtered counts file
Counts <- counts[,select]

# New phenofile with only the variables of interest
Targets <- targets[select,]

# DE-analysis with DESeq2

# Setup design matrix for DE-analysis
condition <- as.character (Targets$Type) # condition of interest to be tested
batch <- as.character (Targets$NGS) # batch effect
sex <- as.character(Targets$Sex) # Sex as a covariate
group_levels <- levels(as.factor(condition))
design <- data.frame(condition=as.factor(condition), batch=batch, sex=sex)
rownames(design) <- colnames(Counts)
dds <- DESeqDataSetFromMatrix(countData=Counts, colData=design, design = ~ batch + sex + condition)

# DESeq2 DE-analysis of the condition of interest, batch effect taken into account, sex as a covariate
dds <- DESeq(dds)

# Display results
res <- results(dds, alpha=0.05)
resOrdered <- res[order(res$padj),]
summary(res)
```

```
## 
## out of 228 with nonzero total read count
## adjusted p-value < 0.05
## LFC > 0 (up)       : 0, 0%
## LFC < 0 (down)     : 0, 0%
## outliers [1]       : 0, 0%
## low counts [2]     : 0, 0%
## (mean count < 8)
## [1] see 'cooksCutoff' argument of ?results
## [2] see 'independentFiltering' argument of ?results
```

```
resOrdered
```

```
## log2 fold change (MLE): condition LS vs CTRL 
## Wald test p-value: condition LS vs CTRL 
## DataFrame with 228 rows and 6 columns
##                         baseMean       log2FoldChange             lfcSE
##                        <numeric>            <numeric>         <numeric>
## hsa-mir-10b-5p  359.127945286884    0.687931059634368 0.237484888316919
## hsa-mir-125a-5p 780.501228445465    0.603420132306963 0.232331286457554
## hsa-mir-127-3p  45.1339945221773    -1.64300477204052 0.627080734315158
## hsa-mir-144-3p  1100.82460008364   -0.959114971832942 0.330034076855235
## hsa-mir-16-5p   371192.343384015    0.863315715176316 0.277753658945226
## ...                          ...                  ...               ...
## hsa-mir-96-5p    93.616441922745   0.0274689378713257 0.456928125815926
## hsa-mir-532-3p  25.1715112485986   0.0365344134704773 0.647935964977894
## hsa-mir-3529-3p 78.4982963166899   0.0126012560579685 0.352736564672473
## hsa-mir-28-3p   200.218788935536 -0.00291543605980014  0.34013614836977
## hsa-mir-425-5p  2271.73513458419  0.00221401883989526 0.185944771023344
##                                 stat              pvalue              padj
##                            <numeric>           <numeric>         <numeric>
## hsa-mir-10b-5p       2.8967361439703 0.00377066742205565 0.267832199418093
## hsa-mir-125a-5p     2.59724009412398 0.00939762103221378 0.267832199418093
## hsa-mir-127-3p     -2.62008491432106 0.00879078748750613 0.267832199418093
## hsa-mir-144-3p     -2.90610891145536 0.00365954048266793 0.267832199418093
## hsa-mir-16-5p       3.10820645335428 0.00188226529460209 0.267832199418093
## ...                              ...                 ...               ...
## hsa-mir-96-5p     0.0601165398218266    0.95206281691499 0.967768229771327
## hsa-mir-532-3p    0.0563858397206331   0.955034437274336 0.967768229771327
## hsa-mir-3529-3p   0.0357242693840633   0.971502218730709 0.980099583498238
## hsa-mir-28-3p   -0.00857137964833629   0.993161112254442 0.993161112254442
## hsa-mir-425-5p    0.0119068626007091   0.990499922640165 0.993161112254442
```

```
mcols(res)$description
```

```
## [1] "mean of normalized counts for all samples"   
## [2] "log2 fold change (MLE): condition LS vs CTRL"
## [3] "standard error: condition LS vs CTRL"        
## [4] "Wald statistic: condition LS vs CTRL"        
## [5] "Wald test p-value: condition LS vs CTRL"     
## [6] "BH adjusted p-values"
```

```
# Create a data frame of the ordered results (top 20)
subset <- head(resOrdered, 20) %>%
  as_tibble(rownames = "miR")

# Create a gene table of the subset
#names(subset)<- c("miR","Mean", "log2FC", "SE", "Wald", "p-value", "FDR")
gt(subset)
```

| miR | baseMean | log2FoldChange | lfcSE | stat | pvalue | padj |
| --- | --- | --- | --- | --- | --- | --- |
| hsa-mir-10b-5p | 359.12795 | 0.6879311 | 0.2374849 | 2.896736 | 0.003770667 | 0.2678322 |
| hsa-mir-125a-5p | 780.50123 | 0.6034201 | 0.2323313 | 2.597240 | 0.009397621 | 0.2678322 |
| hsa-mir-127-3p | 45.13399 | -1.6430048 | 0.6270807 | -2.620085 | 0.008790787 | 0.2678322 |
| hsa-mir-144-3p | 1100.82460 | -0.9591150 | 0.3300341 | -2.906109 | 0.003659540 | 0.2678322 |
| hsa-mir-16-5p | 371192.34338 | 0.8633157 | 0.2777537 | 3.108206 | 0.001882265 | 0.2678322 |
| hsa-mir-32-5p | 438.89304 | -0.7494606 | 0.2812993 | -2.664282 | 0.007715284 | 0.2678322 |
| hsa-mir-361-3p | 147.19465 | 0.9126396 | 0.3363962 | 2.712990 | 0.006667912 | 0.2678322 |
| hsa-mir-423-3p | 720.25553 | -0.6033016 | 0.2136445 | -2.823857 | 0.004744948 | 0.2678322 |
| hsa-mir-15b-5p | 402.36018 | -0.7602167 | 0.3022087 | -2.515535 | 0.011885187 | 0.2709823 |
| hsa-mir-374a-5p | 638.08518 | -1.0141426 | 0.4004843 | -2.532290 | 0.011332013 | 0.2709823 |
| hsa-mir-181d-5p | 40.64623 | -1.2572744 | 0.5091163 | -2.469523 | 0.013529335 | 0.2804262 |
| hsa-mir-15a-5p | 923.51193 | -0.7358426 | 0.3083407 | -2.386460 | 0.017011469 | 0.3216505 |
| hsa-mir-190a-5p | 364.32133 | -0.6953374 | 0.3025734 | -2.298078 | 0.021557323 | 0.3216505 |
| hsa-mir-320a-3p | 1690.77095 | -0.6491387 | 0.2903951 | -2.235364 | 0.025393458 | 0.3216505 |
| hsa-mir-370-3p | 48.76408 | -1.2871668 | 0.5663018 | -2.272934 | 0.023030145 | 0.3216505 |
| hsa-mir-432-5p | 561.06664 | -0.9295053 | 0.4108023 | -2.262659 | 0.023656737 | 0.3216505 |
| hsa-mir-451a | 37683.76940 | -0.7295351 | 0.3252463 | -2.243024 | 0.024895286 | 0.3216505 |
| hsa-mir-654-3p | 63.63660 | -1.4009196 | 0.6009345 | -2.331235 | 0.019740962 | 0.3216505 |
| hsa-mir-107 | 229.77083 | -0.6153741 | 0.2881726 | -2.135436 | 0.032725443 | 0.3391546 |
| hsa-mir-221-3p | 2164.73228 | -0.3891702 | 0.1820512 | -2.137697 | 0.032541366 | 0.3391546 |

```
print("Step 10 completed! There was no difference in c-miR expression between path_MMR carriers with cancer and CTRL group.")
```

```
## [1] "Step 10 completed! There was no difference in c-miR expression between path_MMR carriers with cancer and CTRL group."
```

### 6 Session info

The output from running ‘sessionInfo’ is shown below and details all packages and version necessary to reproduce the results in this report.

```
sessionInfo()
```

```
## R version 3.6.3 (2020-02-29)
## Platform: x86_64-apple-darwin15.6.0 (64-bit)
## Running under: macOS  10.16
## 
## Matrix products: default
## BLAS:   /Library/Frameworks/R.framework/Versions/3.6/Resources/lib/libRblas.0.dylib
## LAPACK: /Library/Frameworks/R.framework/Versions/3.6/Resources/lib/libRlapack.dylib
## 
## locale:
## [1] fi_FI.UTF-8/fi_FI.UTF-8/fi_FI.UTF-8/C/fi_FI.UTF-8/fi_FI.UTF-8
## 
## attached base packages:
## [1] parallel  stats4    stats     graphics  grDevices utils     datasets 
## [8] methods   base     
## 
## other attached packages:
##  [1] DT_0.18                     plotly_4.9.3               
##  [3] DESeq2_1.26.0               SummarizedExperiment_1.16.1
##  [5] DelayedArray_0.12.3         BiocParallel_1.20.1        
##  [7] matrixStats_0.58.0          Biobase_2.46.0             
##  [9] GenomicRanges_1.38.0        GenomeInfoDb_1.22.1        
## [11] IRanges_2.20.2              S4Vectors_0.24.4           
## [13] BiocGenerics_0.32.0         gt_0.2.2                   
## [15] forcats_0.5.1               stringr_1.4.0              
## [17] dplyr_1.0.6                 purrr_0.3.4                
## [19] readr_1.4.0                 tidyr_1.1.3                
## [21] tibble_3.1.1                ggplot2_3.3.3              
## [23] tidyverse_1.3.1             edgeR_3.28.1               
## [25] limma_3.42.2                knitr_1.33                 
## [27] tinytex_0.31                rmarkdown_2.8              
## 
## loaded via a namespace (and not attached):
##   [1] colorspace_2.0-1       ellipsis_0.3.2         htmlTable_2.1.0       
##   [4] XVector_0.26.0         base64enc_0.1-3        fs_1.5.0              
##   [7] rstudioapi_0.13        bit64_4.0.5            AnnotationDbi_1.48.0  
##  [10] fansi_0.4.2            lubridate_1.7.10       xml2_1.3.2            
##  [13] splines_3.6.3          cachem_1.0.4           geneplotter_1.64.0    
##  [16] Formula_1.2-4          jsonlite_1.7.2         broom_0.7.6           
##  [19] annotate_1.64.0        cluster_2.1.2          dbplyr_2.1.1          
##  [22] png_0.1-7              shiny_1.6.0            compiler_3.6.3        
##  [25] httr_1.4.2             backports_1.2.1        lazyeval_0.2.2        
##  [28] fastmap_1.1.0          assertthat_0.2.1       Matrix_1.3-3          
##  [31] cli_2.5.0              later_1.2.0            htmltools_0.5.1.1     
##  [34] tools_3.6.3            gtable_0.3.0           glue_1.4.2            
##  [37] GenomeInfoDbData_1.2.2 rappdirs_0.3.3         Rcpp_1.0.6            
##  [40] cellranger_1.1.0       jquerylib_0.1.4        vctrs_0.3.8           
##  [43] crosstalk_1.1.1        xfun_0.22              rvest_1.0.0           
##  [46] mime_0.10              lifecycle_1.0.0        XML_3.99-0.3          
##  [49] zlibbioc_1.32.0        scales_1.1.1           promises_1.2.0.1      
##  [52] hms_1.0.0              RColorBrewer_1.1-2     yaml_2.2.1            
##  [55] memoise_2.0.0          gridExtra_2.3          sass_0.3.1            
##  [58] rpart_4.1-15           latticeExtra_0.6-29    stringi_1.6.1         
##  [61] RSQLite_2.2.7          genefilter_1.68.0      checkmate_2.0.0       
##  [64] rlang_0.4.11           pkgconfig_2.0.3        bitops_1.0-7          
##  [67] evaluate_0.14          lattice_0.20-44        labeling_0.4.2        
##  [70] htmlwidgets_1.5.3      bit_4.0.4              tidyselect_1.1.1      
##  [73] magrittr_2.0.1         R6_2.5.0               generics_0.1.0        
##  [76] Hmisc_4.5-0            DBI_1.1.1              pillar_1.6.0          
##  [79] haven_2.4.1            foreign_0.8-75         withr_2.4.2           
##  [82] survival_3.2-11        RCurl_1.98-1.3         nnet_7.3-16           
##  [85] modelr_0.1.8           crayon_1.4.1           utf8_1.2.1            
##  [88] jpeg_0.1-8.1           locfit_1.5-9.4         grid_3.6.3            
##  [91] readxl_1.3.1           data.table_1.14.0      blob_1.2.1            
##  [94] reprex_2.0.0           digest_0.6.27          xtable_1.8-6          
##  [97] httpuv_1.6.1           munsell_0.5.0          viridisLite_0.4.0     
## [100] bslib_0.2.4
```
